# The ERK1/2-Elk1, JNK-cJun, and JAK-STAT Transcriptional Axes as Potential Bortezomib Resistance Mediators in Prostate Cancer

**DOI:** 10.1101/2024.04.15.589569

**Authors:** Georgios Kalampounias, Kalliopi Zafeiropoulou, Theodosia Androutsopoulou, Spyridon Alexis, Argiris Symeonidis, Panagiotis Katsoris

## Abstract

The effectiveness of proteasome inhibitors against solid tumors is limited as the emergence of resistance is rapid. Although many mechanisms have been proposed and verified, no definite answer has been given, highlighting the complexity of the resistant phenotype. In this study, a Bortezomib-resistant prostate cancer cell line is created, and a broad-spectrum signaling pathway analysis is performed to identify differences and adaptations the resistant cells exhibit. Our findings highlight the upregulation and activation of Nf-κB, STAT3, cJun, and Elk1 transcription factors in the resistant cells and the subsequent evasion of apoptosis and induction of autophagy, which is constantly activated and substitutes the role of the ubiquitin-proteasome system (UPS). Additionally, assessment of the intracellular reactive oxygen species in resistant cells confirms their downregulation, which is theorized to be a consequence of metabolic changes, increased autophagic flux, and antioxidative enzyme action. The results of this study highlight the potential therapeutic targeting of key kinases and transcription factors, participating in the main signaling pathways and gene regulation of Bortezomib-resistant cells, that could re-sensitize the cells to proteasome inhibitors, thus surpassing the current limitations.

## 1. Introduction

Proteasome inhibitors were first used as anticancer agents more than 20 years ago and have since saved thousands of lives from multiple myeloma, mantle cell lymphoma, and many other hematological malignancies (1). They target the proteasome, a protein multi-catalytic complex that actively participates in the cell’s homeostatic mechanisms by selectively degrading poly-ubiquitinated polypeptides. The proteasome is of paramount importance in cancer cells, and its increased activity and subunit accumulation are well-documented in various cancer types. It is believed that cancer cells, due to increased biosynthetic rates and miss-regulated control mechanisms, produce more miss-folded polypeptides that need to be recycled (2,3). Additionally, many proteins inside the cell have very specific turnover rates that are regulated by proteasomal degradation through their labeling with ubiquitin chains, and therefore, impairment of their lifecycle can affect cellular functions like the progression of the cell cycle, mitochondrial function, and gene expression (4,5). Cancer cells have been found to rely on the proteasome to “tune” this system, and this mechanism, namely the ubiquitin-proteasome system (UPS), is a very important pathway that, once targeted and sabotaged, can induce apoptosis and lead to cell death (6).

Since their discovery, proteasome inhibitors have been tested on various cell lines, animal models, and even clinical studies; however, besides a group of hematological cancers, they are not very efficient against solid tumors, as resistance emerges (7–15). Bortezomib was the first proteasome inhibitor used for its tumoricidal properties and acts by binding to the main catalytic subunit of the 20S proteasome, the β5 subunit, which has chymotrypsin-like activity. Patients who received Bortezomib as treatment and relapsed due to resistance were found to have mutations in the *PSMB5* gene, the one coding for the β5 subunit, that significantly reduced the drug binding ability or caused increased expression (16–21). However, those mutations are not the sole ways for resistance to emerge, since most of the relapsed patients did not have alterations regarding β5 structure or expression. The current hypothesis is that, besides genetic mutations that may alter the proteasome subunits’ abundance and structure, thus rendering Bortezomib insufficient, a great role in the resistant phenotype is also played by changes in the signaling cascades that regulate apoptosis, autophagy, and oxidative stress (22–27).

Nf-κB is a transcription factor of great interest in immunology; however, it was one of the first molecules to be found active in Bortezomib-resistant cells, implying a role in the emergence of that aggressive phenotype (28). The STAT family is another class of transcription factors that have been found active in resistant clones, as well as the ERK1/2 signaling pathway that mostly regulates cell survival and proliferation (29–34). Recent advances in Bortezomib resistance, focused on prostate cancer, report the activation of autophagy in the resistant cells and describe a regulation mechanism that substitutes the impaired UPS and abolishes the cells’ dependence on it to maintain their proteostasis (27,35–38). Though Beclin-1 regulates autophagic flux to successfully downregulate pro-apoptotic pathways and diminish oxidative stress levels, given the complexity of resistance emergence and the multiple layers that a resistant phenotype can exhibit, the need for a new approach is of paramount importance. This study aims to investigate the phenotype of a prostate cancer cell line that acquires resistance to Bortezomib and monitor how the cells’ signaling pathways regulating main biological functions have adapted, possibly suggesting new pharmaceutical targets. Focusing on the new players in this chess game—autophagy and oxidative stress—and unraveling and targeting their regulation mechanisms could renew interest in proteasome inhibitor therapy, thus better armoring us against drug resistance.

## 2. Results

### 2.1. Creation of the PC-3 RB40 Cell Line and Monitoring Main Biological Functions

#### 2.1.1. IC_50_ Determination and Proliferation Assays

After long-term exposure to increasing concentrations of Bortezomib, the PC-3 cell line gradually developed resistance to the drug. During the dose escalation, the cell morphology was altered, and the cells were used to create long protrusions instead of maintaining the typical PC-3 morphology. This phenomenon was evident in the naïve cells and persisted in the resistant clone as well during the first three weeks of stable Bortezomib administration (40 nM). This was theorized as a sign of poor adaptation and stress, and the signaling assays were not conducted until the cell morphology was restored. During this interval, proliferation assays were performed, and the IC_50_ of the resistant cells was just above the stable medium concentration of 40 nM (Table 1) (Figure A1). Following a total of 32 weeks, the cell morphology was restored, and the resistant cell clone had elevated its resistance capacity from 40 nM to about 55 nM, as shown by their IC_50_ values (Figure 1a). Since the 40 nM dose was the lowest concentration where no naïve PC-3 cells survived after 72 h of treatment while the resistant cells were adequately adapted, it was established as the main context for signal transduction, apoptosis, and stress assays. This cell clone of PC-3 was named PC-3 RB40 and was constantly cultured in RPMI 1640 medium supplemented with 40 nM Bortezomib. At this time point/resistance capacity, the cells were also assessed for resistance against the second-generation proteasome inhibitor Carfilzomib (Figure 1b) as well as the anthracycline Doxorubicin, which is a known tumoricidal agent known to induce apoptosis (Figure 1c). Normally, PC-3 cells are susceptible to Doxorubicin treatment, which has a 48-hour IC_50_ value of 35 nM (Table 1). Our experiments showed that the RB40 cell clone had the same IC_50_ as the naïve cells, indicating that no multidrug resistance mechanism had emerged but rather specific resistance to proteasome inhibitors (Table 1). The RB40 clone also exhibited cross-resistance to Carfilzomib; however, the IC_50_ value was significantly lower. These results were confirmatory of a previous study conducted in our laboratory, where DU-145 cells were used as a model for Bortezomib resistance, and the resulting phenotype was a cell line resistant only against proteasome inhibitors (Bortezomib and Carfilzomib) and not against anthracyclines (27).

**Figure 1.**
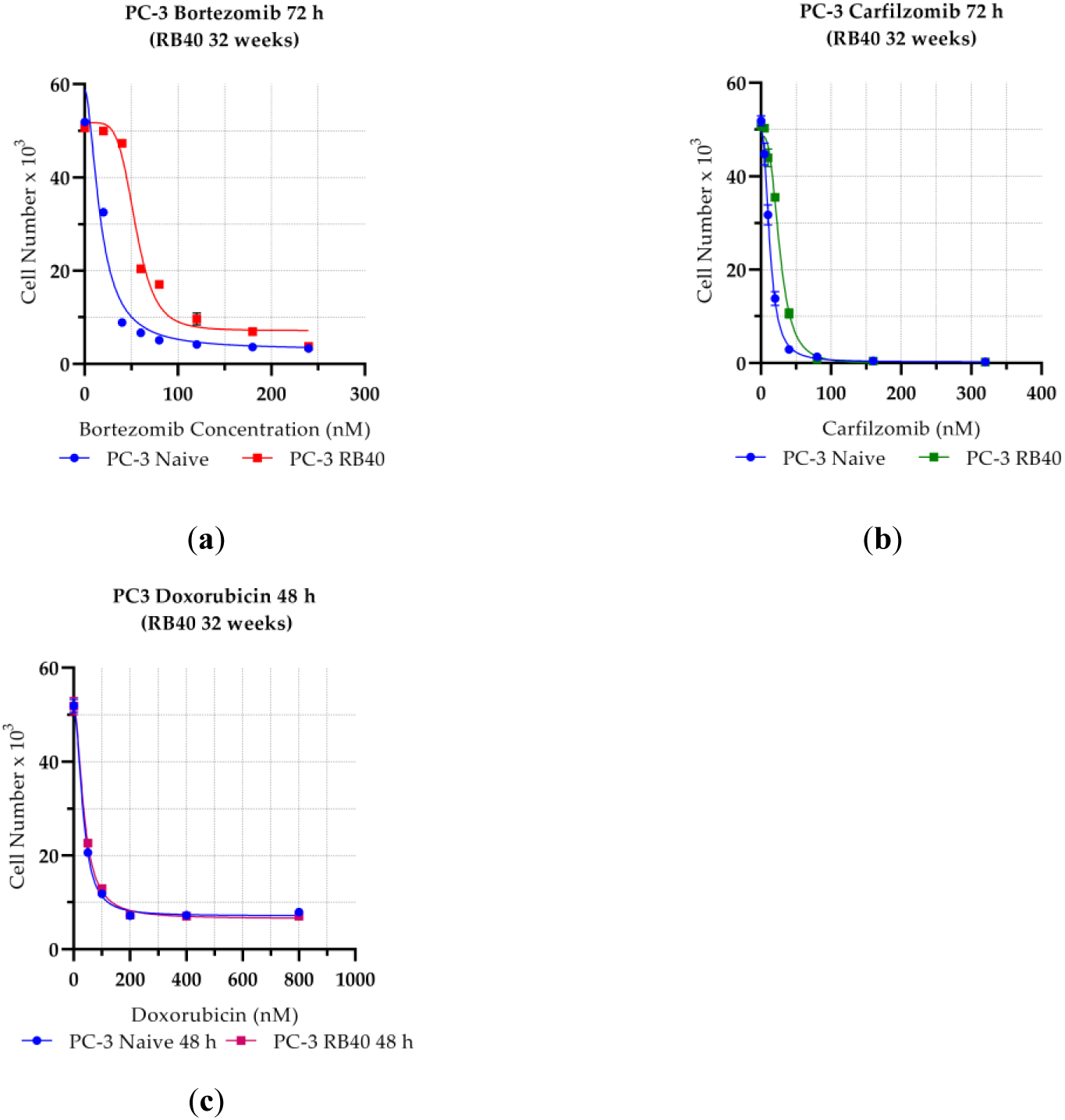
Assaying the Effects of Bortezomib, Carfilzomib, and Doxorubicin on the Proliferation of Naïve PC-3 and PC-3 RB40 Cells. Equal numbers of cells were cultured for 72 h in 24-well microplates with various concentrations of (**a**) Bortezomib (0-240 nM); (**b**) Carfilzomib (0-325 nM); and (**c**) Doxorubicin (0-1000 nΜ), and the live cells were measured using the Crystal Violet Assay. Each dot represents the average of three experimental values, and the error bars represent the standard error of the mean (SEM). The fitting line was graphed in Prism 8 using the built-in model for IC50 determination. The blue lines represent the proliferation curves of naïve PC-3 cells, the red line the proliferation curve of PC-3 RB40 cells during Bortezomib treatment, the green line the curve of PC-3 RB40 cells during Doxorubicin treatment. Each plot represents one experiment, while the mean IC50 of three replicated experiments is presented in Table 1.

**Table 1.** The IC50 values of naïve and resistant cells following 72 h of Bortezomib/Carfilzomib or 48 h of Doxorubicin treatment. Each value represents the average of three replicate experiments, and the annotated error is the standard error of the mean (SEM). The data were analyzed in Prism 8 using the built-in tools for IC50 determination.

| Drug<br>(incubation time) | Time elapsed<br>(weeks) | IC<sub>50</sub> (nM) |  | IC<sub>50</sub> ratio<br>(resistant: naïve) |
| --- | --- | --- | --- | --- |
|  |  | Naïve PC-3 | PC-3 RB40 |  |
| Bortezomib (72 h) | 4 | 15.07±2.14 | 21.87±1.72 | 1.45 |
|  | 12 | 17.30±2.03 | 25.47±0.89 | 1.47 |
|  | 24 | 16.39±1.06 | 40.23±0.98 | 2.45 |
|  | 28 | 16.12±0.11 | 48.56±1.11 | 3.01 |
|  | 32 | 16.96±1.1 | 54.64±1.65 | 3.22 |
| Carfilzomib (72h) | 4 | 12.86±1.34 | 13.78±2.01 | 1.07 |
|  | 32 | 13.11±0.45 | 24.17±2.04 | 1.84 |
| Doxorubicin (48 h) | 4 | 34.18±4.12 | 36.81±2.62 | 1.08 |
|  | 32 | 33.23±3.17 | 35.19±3.01 | 1.06 |

#### 2.1.2. The Resistant Cells Fully Restore their Cell Cycle Progression Rate and Successfully Evade Apoptosis

To validate the resistant cells’ adaptation to the proteasome inhibitor-rich medium, we assessed their cell cycle progression and their apoptosis rate. Bortezomib has been demonstrated to cause both G_1_/S arrest as well as G_2_/M arrest, and this mechanism also triggers apoptosis (39–41). Naïve PC-3 cells, after 24 hours of treatment with 40 nM Bortezomib, exhibited the bibliographically reported G_2_/S arrest (and, to a lesser extent, G_1_/S arrest), which is shown as an accumulation of cells in the second peak (corresponding to G_2_-and M-phase DNA content) when the cells were stained with propidium iodide (Figures 2a, 2b). At the same dose, the PC-3 RB40 cells indicate only a mild cell cycle distortion (Figure 2d), and they even maintain (untreated) naïve-like cell cycle progression even after 40 nM of Bortezomib (data not shown). The untreated PC-3 RB40 cells were documented to have a naïve-like distribution among the three cell cycle phases (Figure 2c). These results were by the time needed for the cells to fully form a monolayer inside cell culture dishes, which was the same for untreated naïve PC-3 cells and PC-3 RB40 cells.

**Figure 2.**
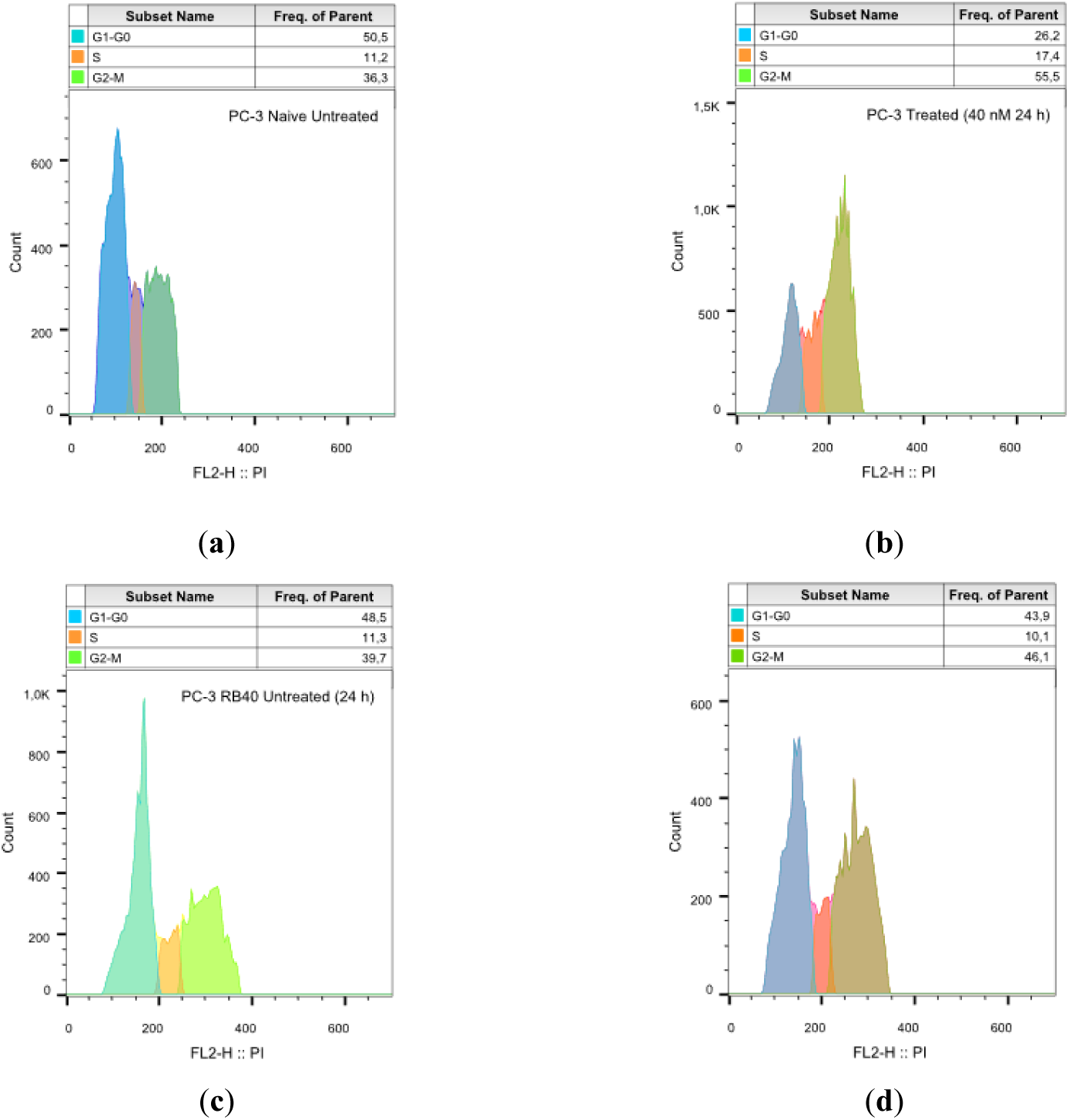
Cell Cycle Analysis of Naïve PC-3 and PC-3 RB40 Cells Following Treatment with Bortezomib. Equal numbers of cells were cultured inside 100-mm dishes, and 24 h before analysis, the media were replaced. (**a**) Naïve PC-3 cells were cultured for 24 h in RPMI 1640 medium supplemented with 10% FBS (experimental control of baseline cell cycle progression); (**b**) Naïve PC-3 cells were cultured in medium containing 40 nM of Bortezomib for 24 h; (**c**) PC-3 RB40 cells were cultured for 24 h in RPMI 1640 medium supplemented with 10% FBS and; (**d**) PC-3 RB40 cells were cultured for 24 h in RPMI 1640 medium supplemented with 10% FBS and 40 nM of Bortezomib (basal culture conditions for this cell line). The cells were fixed and permeabilized with methanol, treated with RNAase, and stained with propidium iodide. Equal numbers of events were acquired using a FACS Calibur flow cytometer by measuring the propidium iodide, which is correlated to the DNA content, so the different cell cycle phases (G1/G0, S, and G2/M) could be shown in histograms using the FlowJo software. The figure presents a representative experiment. The same procedure was replicated three times.

The cells were also assessed for their apoptosis rate. An Annexin-V/PI kit was used that stains phosphatidylserine residues to indicate their extracellular presence (early apoptotic marker) and the membrane’s integrity, which is a marker of late apoptosis or necrotic/ferroptotic cell death. The cells that were cultured with increasing doses of Bortezomib were initially susceptible to apoptosis induction, despite the constant drug presence; however, following eight weeks of the stable, high Bortezomib dose of 40 nM, they had fully adapted. This was evident after assessing them for apoptosis, as the apoptotic activity of PC-3 RB40 cells (cultured with Bortezomib) had dropped to levels comparable to those of naïve cells (Figures 3a, 3c-3e) while the treated cell population of naïve cells indicated high levels of apoptotic death (Figure 3b). Further dose escalation on the resistant clone (up to 80 nM) significantly induced apoptosis; however, even this high Bortezomib dose (Figure 3f) did not cause cell death comparable to that of naïve cells (Figure 3b).

**Figure 3.**
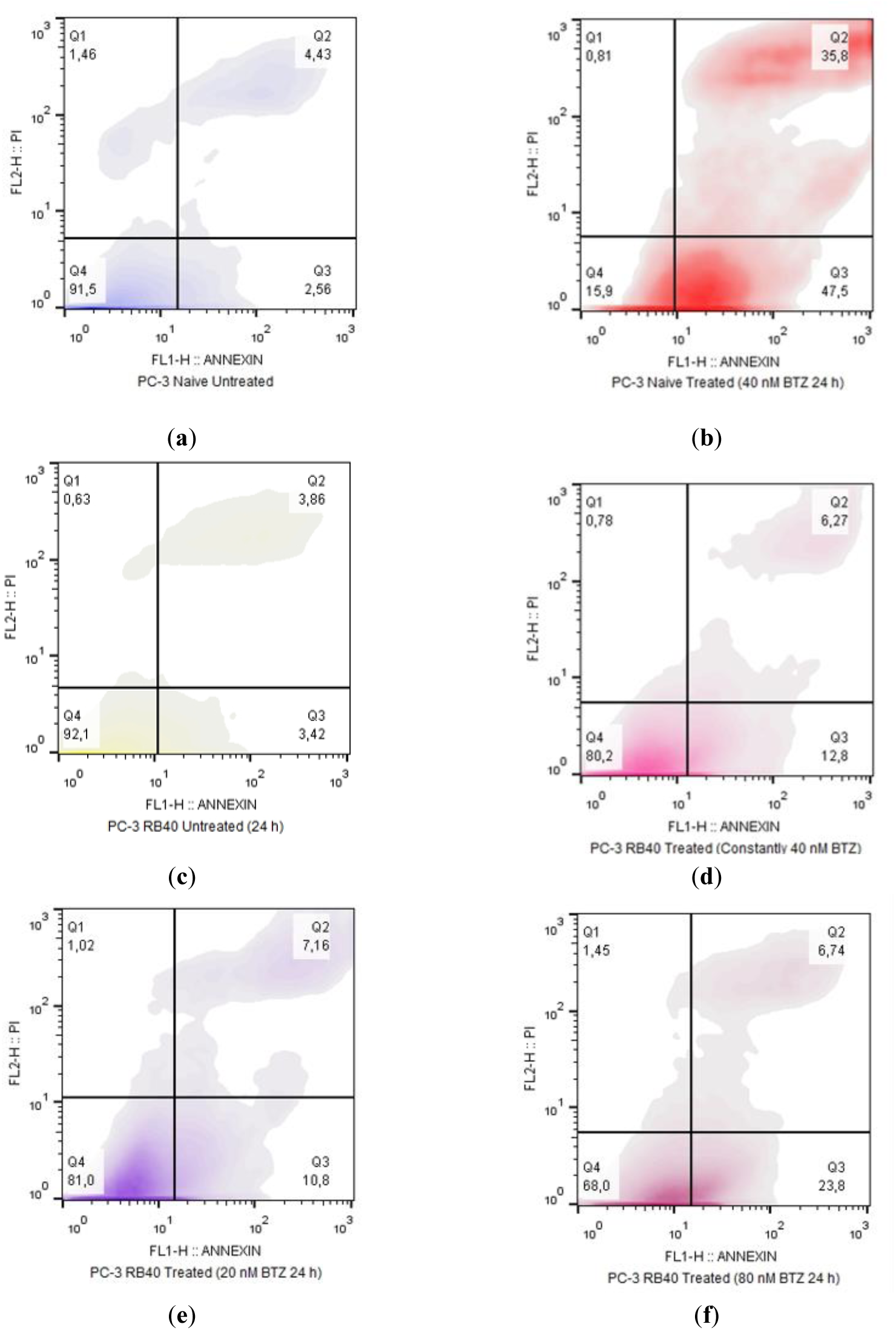
Apoptosis Assay of Naïve PC-3 and PC-3 RB40 Cells Following Treatment with Bortezomib. Equal numbers of cells were cultured inside 100-mm dishes, and 24 h before analysis, the media were replaced. (**a**) Naïve PC-3 cells were cultured for 24 h in RPMI 1640 medium supplemented with 10% FBS (blue coloring)(experimental control of baseline cell cycle progression); (**b**) naïve PC-3 cells were cultured in medium containing 40 nM of Bortezomib for 24 h (red coloring); (**c**) PC-3 RB40 cells were cultured for 24 h in RPMI 1640 medium supplemented with 10% FBS (yellow coloring); (**d**) PC-3 RB40 cells were cultured for 24 h in RPMI 1640 medium supplemented with 10% FBS and 40 nM of Bortezomib (magenta coloring)(basal culture conditions for this cell line); (**e**) PC-3 RB40 cells were cultured for 24 h in RPMI 1640 medium supplemented with 10% FBS and 20 nM of Bortezomib (purple coloring); and (**f**) PC-3 RB40 cells were cultured for 24 h in RPMI 1640 medium supplemented with 10% FBS and 80 nM of Bortezomib (lilac coloring) The cells were stained with an Annexin-V/propidium iodide (PI) kit inside a Calcium-containing buffer to measure the levels of apoptosis. Equal numbers of events were acquired using a FACS Calibur flow cytometer by measuring the fluorescence emitted by FITC, which is bound to Annexin-V and PI, and the data were analyzed using the FlowJo software. The graphs display the density plots of Annexin-V and PI, and the four different quartiles correspond to Q1: necrotic cells; Q2: cells in late apoptosis; Q3: cells in early apoptosis; and Q4: live cells. The figure presents a representative experiment. The same procedure was replicated three times.

#### 2.1.3. Bortezomib Fails to Inhibit Migration of Resistant Cell, Epithelial to Mesenchymal Transition (EMT) Markers are Expressed and the Wnt Signaling Pathway is Active

Besides the known effects on cell proliferation and apoptosis, Bortezomib has been shown to inhibit cell adhesion and migration by interfering with the turnover times of various molecules and thus disrupting the function of the adhesomes (42–44). Besides turnover dysregulation, ubiquitin also serves as a signaling molecule, controlling Wnt signaling through stabilization or selective degradation; therefore, the consequences of UPS impairment were assessed (45–48).

Naïve and resistant cells (PC-3 RB40) were assessed using the Scratch Test assay to observe differences in the cells’ ability to successfully heal artificial wounds by dividing and migrating. Dose-response experiments showed that Bortezomib inhibited wound healing only in the naïve cells, beginning with concentrations as low as 40 nM (Figures 4a-4c), while the RB40 was not affected by the inhibitor, maintaining its migratory capabilities at concentrations greater than 40 nM (Figures 4c, 4d).

**Figure 4.**
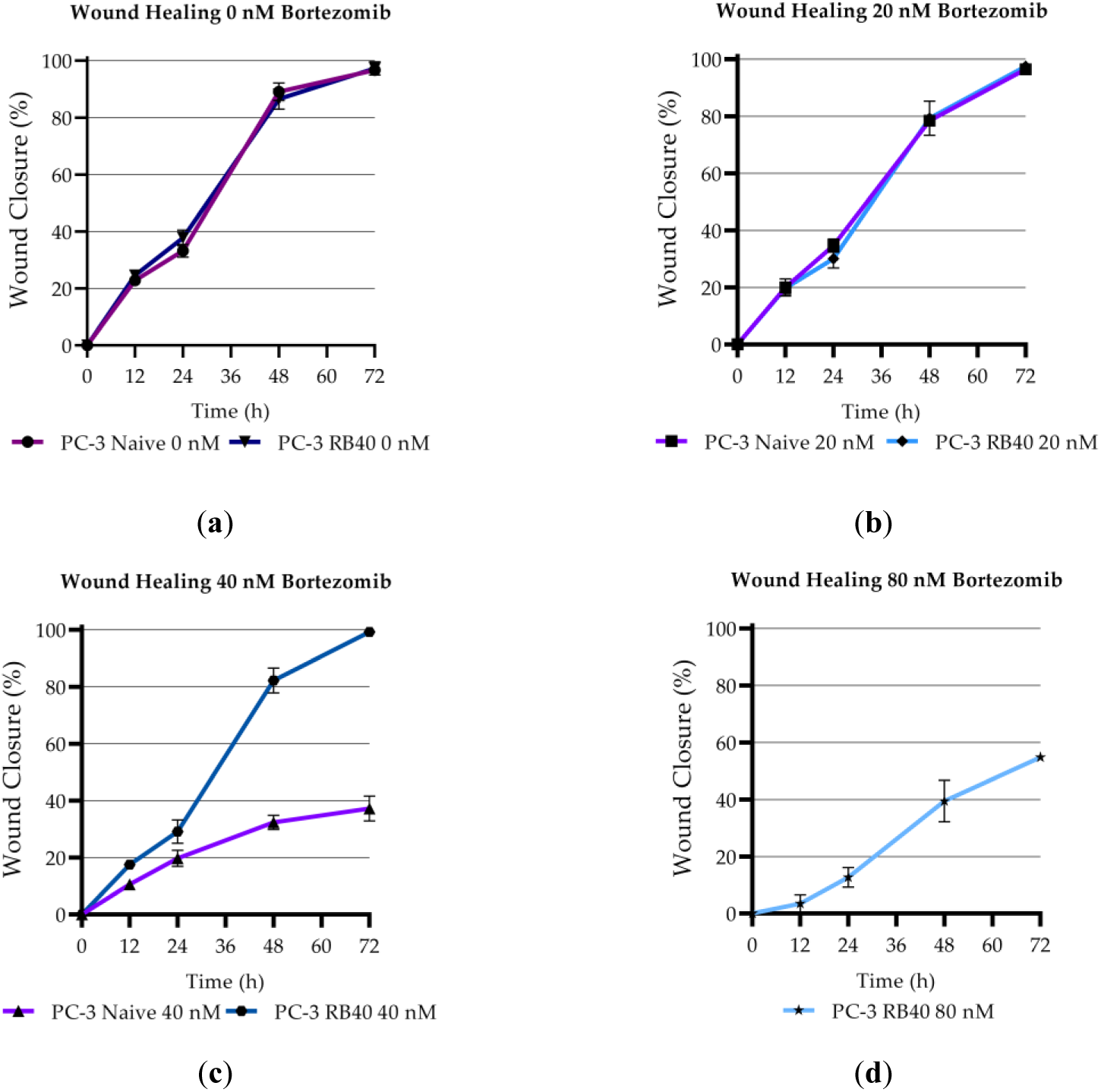
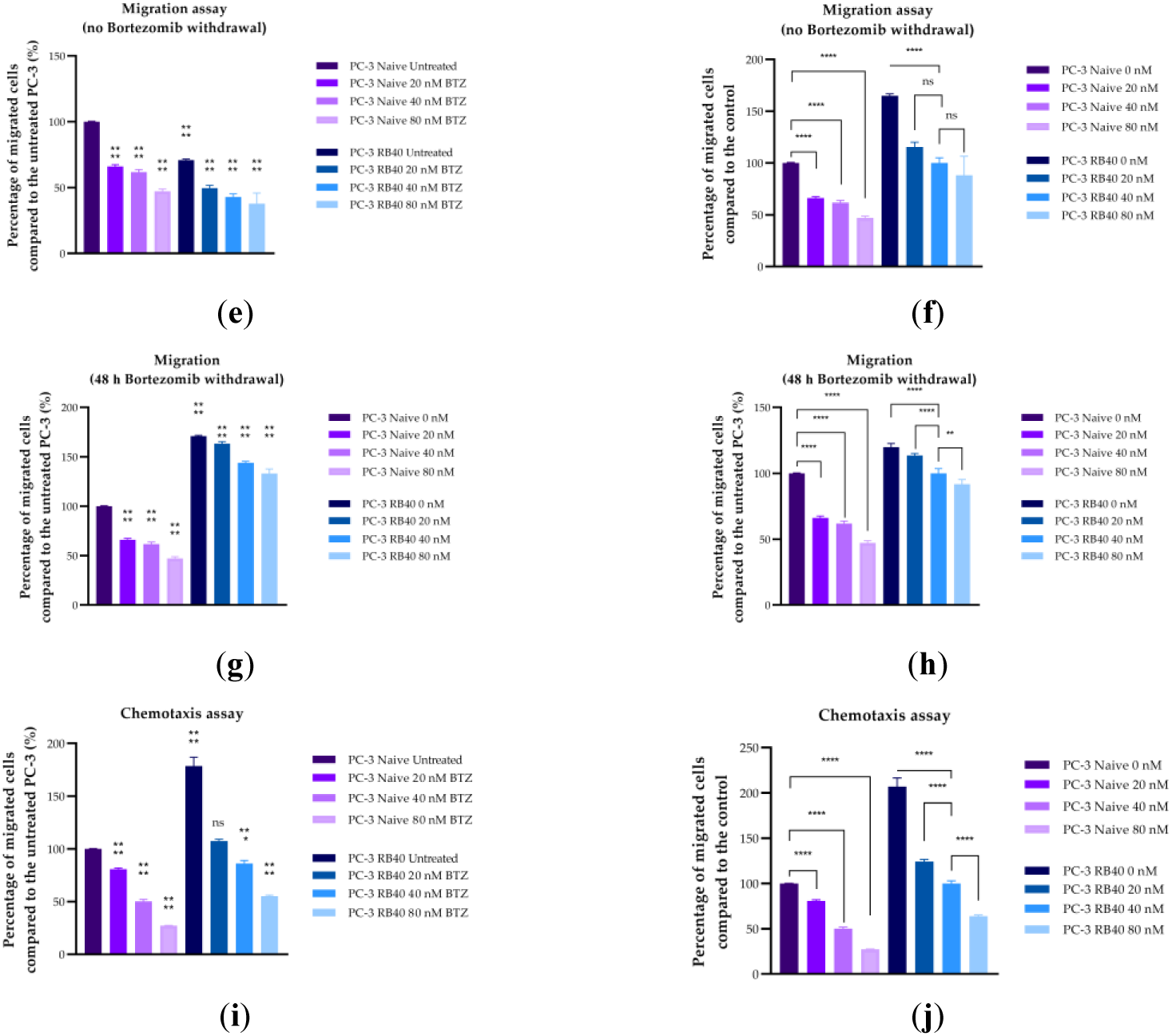
Assaying the Effects of Bortezomib on the Migration of Naïve PC-3 and PC-3 RB40 Cells. (**a-d**) Scratch Test/Wound Healing Assay; Cells were cultured in 6-well plates until confluency, and then scratches were made. Various concentrations of Bortezomib (0, 20, 40, 80 nM) were added, and photographs were taken at the key time points of 0, 24, 48, and 72 h using a camera mounted on an inverted microscope at 100X magnification. The photographs showing wound closure were then analyzed using an ImageJ plug-in, and the healing rates were determined. The data were plotted in Prism 8. (**a**) Naïve PC-3 and PC-3 RB40 cells untreated for 72h; (**b**) Naïve and PC-3 RB40 cells, both treated with 20 nM Bortezomib. (**c**) Naïve and PC-3 RB40 cells, both treated with 40 nM Bortezomib. (**d**) RB40 cells treated with 80 nM Bortezomib. Each wound healing experiment was conducted in triplicate, and the values on the plots are the averages. The error bars correspond to the standard error of the mean (SEM) from the three experiments. (**e, f**) Migration Assay; Equal numbers of cells were placed in Transwell/Boyden Chambers in serum-free RPMI 1640 medium containing increasing concentrations of Bortezomib (0, 20, 40, and 80 nM), and the inserts were placed in microwells containing FBS-supplemented medium. The cells were left for 24 h to migrate, and afterwards, the cells crossing the filters were fixed and stained with crystal violet. Photographs were taken using a 10x objective and the fixed cells were counted using the multipoint tool by ImageJ; (**e**) The percentage of migrated cells were compared to the naïve untreated sample; (**f**) The percentages of total migrated cells, compared to each clone’s baseline conditions (untreated for the naïve cells, and 40 nM Bortezomib for the RB40 cells); (**g, h**) The migration experiments were also conducted following a 48-hour Bortezomib withdrawal/ clearance period of the RB40 clone. (**i, j**) Chemotaxis Assay; Equal numbers of cells were placed in Transwell/Boyden Chambers in serum-free RPMI 1640 medium, and the inserts were placed in microwells with FBS-supplemented medium containing increasing concentrations of Bortezomib (0, 20, 40, and 80 nM). The cells were left for 24 h to migrate, and afterwards, the cells crossing the filters were fixed and stained with crystal violet. Photographs were taken using a 10x objective, and the fixed cells were counted using the multipoint tool by ImageJ; (**i**) The percentage of migrated cells were compared to the naïve untreated sample; (**j**) The percentages of total migrated cells, compared to each clone’s baseline conditions (untreated for the naïve cells, and 40 nM Bortezomib for the RB40 cells). The results were analyzed using multiple comparisons of one-way ANOVA in the Prism 8 software. (* corresponds to P =0,01; ** corresponds to P =0,001; *** corresponds to P =0,0001; and **** corresponds to P <0,0001). The error bars represent the standard error of the mean (SEM).

The assays lasted 72 h, and at this point, the untreated naïve cells as well as the RB40 clone (untreated and treated with 20 or 40 nM Bortezomib) managed to fully heal the scratches. Full images of a representative wound healing experiment can be found in Appendix B (Figure A2).

The migratory and chemorepellent effects of Bortezomib on both naïve and resistant cells were subsequently assessed using Transwell chambers (Figures 4e-4j). Bortezomib exhibited suppressive effects on the cells’ ability to migrate toward the chemoattractant medium, as indicated by experiments where the drug was placed in the insert with a serum-free medium. (Figures 4e, 4f). The effect was also observed in resistant cells whose migratory abilities were severely impaired in the presence of Bortezomib compared to the drug absence. Surprisingly, the resistant cell’s ability to migrate even in drug absence was lower than that of naive cells. However, the ability of untreated RB40 cells to migrate was greater than that of low-dose treated naïve cells. The two conditions may seem different; however, given that the resistant cells were not given a Bortezomib clearance period from their previous maintenance in 40 nM Bortezomib, the decreased levels of migration could be a consequence of the residual drug. Therefore, we repeated the experiment by adding a Bortezomib clearance period of 48 h and the results indicated that the RB40 cells line was significantly more aggressive compared to naïve cells (Figures 4g, 4h). Bortezomib (both in drug-deprived and treated-resistant cells) acted in a dose-dependent manner; however, in the drug-deprived cells’ case, even 80 nM of Bortezomib did not diminish the cell’s migratory potential to naïve levels.

Bortezomib was also examined as a chemoattractant/chemorepellent agent by placing it in the lower compartment (microplate well) along with serum at a 10% concentration (Figures 4i, 4j). It successfully repelled naïve PC-3 cells and masked the chemoattractant medium’s presence; however, the resistant cells were not affected by the drug’s presence at the same levels. Notably, the baseline cell motility levels for both cell clones (intreated naïve cells and resistant cells treated with 40 nM) were almost identical, indicating that the resistant cells can ignore the presence of Bortezomib, successfully migrating towards the chemoattractant medium. Full images of representative migration and chemotaxis assays can be found in Appendix B (Figure A3).

The resistant cells’ ability to defy the anti-migratory effects of Bortezomib regarding cell motility and migration led to the conclusion that proteins related to those functions could have been affected; therefore, basic cadherins, α_ν_β_3_-integrin, and β-catenin were assayed using western blots. N-cadherin and E-cadherin are two of the most important calcium-dependent cell adhesion molecules, both participating in a phenomenon called cadherin-switch, during which N-cadherin is upregulated and E-cadherin is down-regulated, both driving an aggressive phenotype (49,50). The resistant clone emerged, significantly increasing the accumulation of N-cadherin compared to the baseline expression of naive cells (Figures 5a, 5b). Treatment with Bortezomib increased N-cadherin accumulation both on naïve and resistant cells; however, the resistant cells had a more stable pattern of expression, regardless of the drug’s presence. The opposite phenomenon was documented regarding E-cadherin. Following treatment with 20 nM of Bortezomib, E-cadherin indicated reduced accumulation; however, a higher dose of Bortezomib (40 nM) led to a significant decrease (Figures 5a, 5c). In the resistant clone, the cadherin switch was evident since E-cadherin had significantly lowered accumulation levels compared to the (naïve) untreated cells. Additionally, E-cadherin accumulation did not fluctuate in this clone, regardless of the inhibitor dose, showing an abolishment of ubiquitin regulation (Figure 5c).

**Figure 5.**
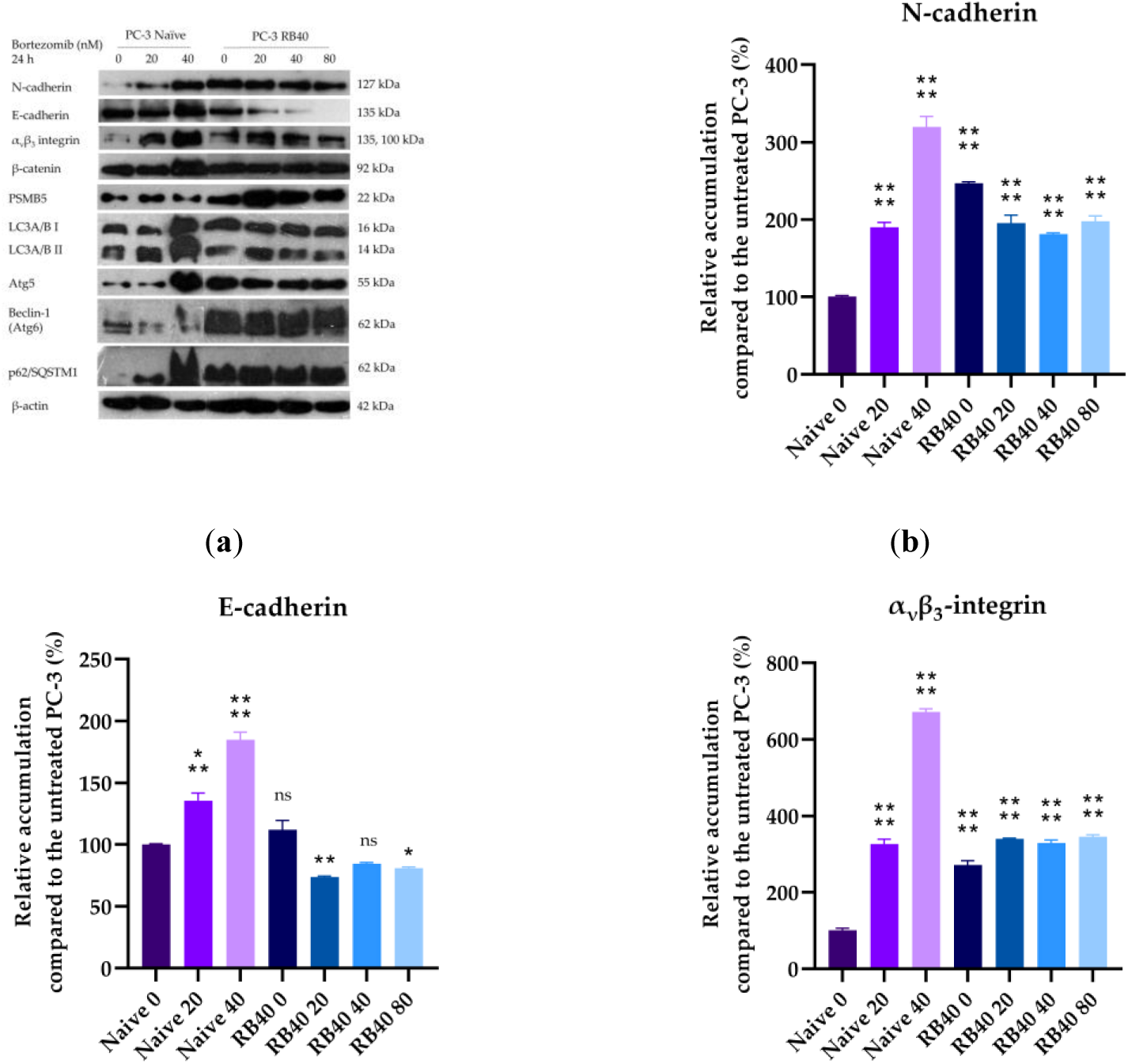
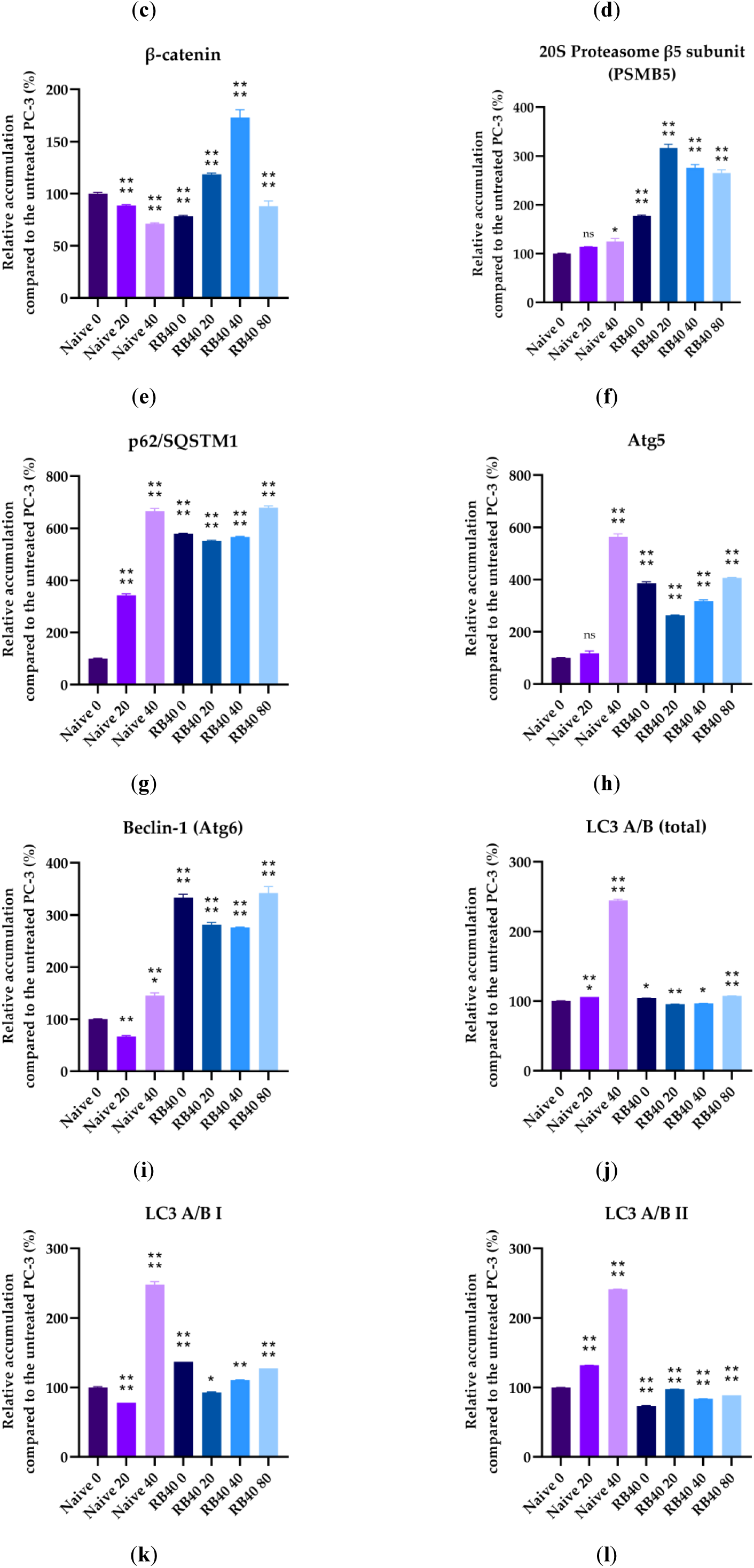
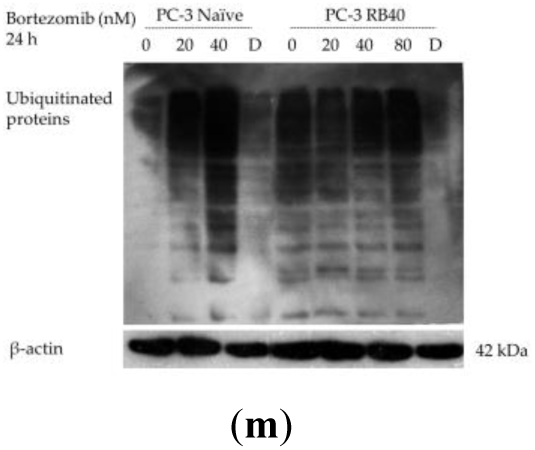
Western Analysis of N-cadherin, E-cadherin, ανβ3-integrin, β-catenin PSMB5, p62/SQSTM1, Atg5, Beclin-1, LC3A/B, and ubiquitinated proteins. Cells (Naïve PC-3 and PC-3 RB40) were cultured inside 100 mm dishes, and 24 h before confluency, the media were changed and fresh RPMI 1640 supplemented with 10% with or without the designated bortezomib doses (20, 40, 80 nM) was added. Following 24 h of incubation, the cells were lysed using RIPA buffer, and equal quantities of total proteins were loaded onto 12% polyacrylamide gels and analyzed with SDS-PAGE. The protein content was determined using the Bradford assay, and as a validation of successful transfer, the total proteins on the gel and the membrane were stained with a Coomassie Brilliant Blue Solution. Western analysis of β-actin was used as a reference protein after it was validated that its accumulation remained stable during all procedures. (**a**) Using specific polyclonal antibodies, the accumulation of N-cadherin, E-cadherin, ανβ3-integrin, β-catenin, PSMB5, p62/SQSTM1, Atg5, Beclin-1 and LC3A/B was detected using the SuperSignal™ West Femto Maximum chemiluminescence Kit. The chemiluminescence was developed on autoradiography films, which were subsequently scanned. Molecular weight markers were used during SDS-PAGE and the approximate molecular weight of each detected polypeptide is annotated next to the band. (**b-l**) Quantification of the scanned blots was followed using the plug-in “Gels” in ImageJ after conversion to grayscale images, and the band intensities and bit depth were calculated. The data were retrieved from triplicate experiments and after normalization using β-actin, bar charts were created. Each bar represents the average relative accumulation of the target protein compared to the untreated naïve PC-3 cells (first blot lane). The results were analyzed using multiple comparisons of one-way ANOVA in the Prism 8 software. (* corresponds to P =0,01; ** corresponds to P =0,001; *** corresponds to P =0,0001; and **** corresponds to P <0,0001). The error bars represent the standard error of the mean (SEM). (**m**) To obtain information about the ubiquitination, a mouse polyclonal antibody was used, and a membrane was appropriately probed. The resulting film was scanned, and the results were presented as they were.

The accumulation of α_ν_β_3_-integrin was also assessed and indicated a dose-dependent accumulation in naïve cells that reached a six-fold increase compared to the control sample (Figures 5a, 5d). The resistant cells maintained higher integrin levels (three-fold), even when the drug was absent, and during Bortezomib treatment (20–80 nM), the protein levels were not affected. α_ν_β_3_-integrin is one of the most studied adhesion molecules in prostate cancer, having been reported as essential for extracellular matrix adhesion during invasion and metastasis (51). α_ν_β_3_-integrin interacts not only with the actin cytoskeleton and the related scaffold proteins but also regulates survival and drives metastasis-related genes by clustering and acting as recruitment areas where phosphorylation of FAK can take place. These signals are finally transmitted inside the nucleus through the ERK1/2 pathway, which in our experiments were found to be activated as will be discussed subsequently. Besides ERK1/2 signaling, α_ν_β_3_-integrin also regulates MMP activity through PI3K signaling (51). Given that PI3K-Akt signaling was found to be upregulated as will be discussed later, the role of α_ν_β_3_-integrin in the resistant cells’ aggressive phenotype regarding migration was adequately explained.

Finally, a connecting link between the UPS and adhesion is also the transcription regulator β-catenin, a molecule that has also been correlated with cancer invasiveness and EMT (52). The resistant clones exhibited increased β-catenin accumulation, an observation in accordance with the protein’s role as an oncogene (53). Following treatment with Bortezomib, the naïve cells decreased the accumulation of the protein; however, in the resistant cells, the baseline levels (at 40 nM of Bortezomib) were increased by 50% compared to the untreated naïve cells. Deviation from this concentration affected β-catenin negatively; however, the total accumulation ranged between values greater than those observed in the naïve clone (Figures 5a, 5e).

### 2.2. Autophagy Substitutes the Impaired Proteasome-Ubiquitin System in Resistant Cells and Oxidative Stress Levels Drop

#### 2.2.1. The Ubiquitination levels of Total Proteins are Restored in the Resistant Clones

The presence of Bortezomib interferes with the UPS, leading to the accumulation of poly-ubiquitinated polypeptides that cannot be degraded through cell mechanisms. Therefore, the accumulation can act as a marker for UPS activity, and after western blot analysis, the impairment of the UPS on the naïve cline was made evident in a dose-dependent manner (Figure 5m). The resistant cell exhibited baseline ubiquitination levels, which could mean either de-sensitization to Bortezomib or exploitation of alternative pathways to substitute for the blocked proteasome. Bortezomib binds to the 20S proteasome’s β5 subunit, and in many Bortezomib-resistance cases, mutations or alterations in the subunit expression have been documented (16,17,54). Additionally, in these experiments, both naïve and resistant cells were incubated with Doxorubicin, which has been found to activate the UPS system (55). Indeed, our experiments indicated decreased levels of ubiquitination, which was theorized as an index of activated proteasome-mediated proteolysis (Figure 5m). The β5 (or PSMB5) subunit was assessed using western blots and found to be elevated in the resistant cells (Figures 5a, 5f). This was theorized as a response to the proteasome’s inability to process the poly-ubiquitinated protein load and was not viewed as a key characteristic of Bortezomib resistance because of the overwhelming Bortezomib concentration that the resistant cells were able to withstand. For this reason, our interest was directed towards the activation of autophagy as a response mechanism.

#### 2.2.2. Bortezomib-Resistant Cells Express Autophagy Markers and Indicate Persistent and Increased Autophagic Flux

The idea that autophagy can substitute for dysregulated UPS has been documented in several studies of different cancer types, including prostate cancer (27,35,56,57). To monitor the autophagic activity, the lysosomal marker Lysotracker RED was used. Lysotracker has been used as an autophagy marker, even though it does not directly assess autophagic flux but rather stains and thus quantifies acidic proteins (58–60). The acidic protein content of a cell is correlated to the lysosomal load, and therefore, due to the organelles’ role in autophagy, Lysotracker can be used to study autophagic activity. Staining with Lysotracker revealed that treatment with Bortezomib significantly increased autophagic flux in both naïve and resistant cells (Figure 6). The resistant cells were documented to possess high autophagy levels even in the absence of the drug compared to the naïve cells (Figures 6a, 6c). Incubation with Bortezomib in both naïve and resistant cells also induces autophagy in a dose-dependent manner (Figures 6b, 6d).

**Figure 6.**
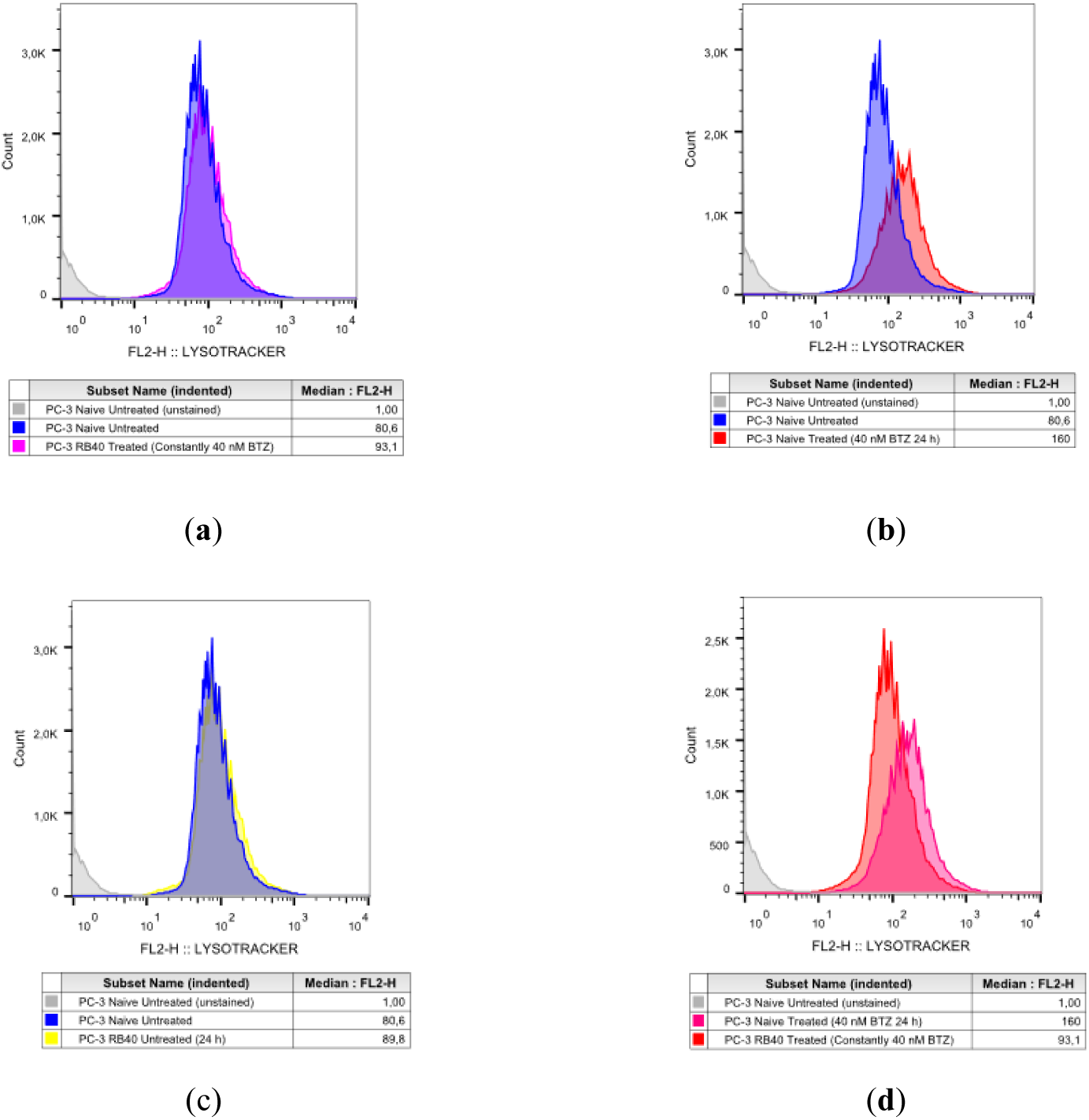
Autophagy Assay of Naïve PC-3 and PC-3 RB40 Cells Following Treatment with Bortezomib. Equal numbers of cells were cultured inside 100 mm dishes, and 24 h before analysis, the media were replaced. (**a**) Naïve PC-3 cells were cultured for 24 h in RPMI 1640 medium supplemented with 10% FBS (blue coloring)(experimental control of baseline cell cycle progression); (**b**) Naïve PC-3 cells were cultured in medium containing 40 nM of Bortezomib for 24 h (magenta coloring); (**c**) PC-3 RB40 cells were cultured for 24 h in RPMI 1640 medium supplemented with 10% FBS and (yellow coloring); (**d**) PC-3 RB40 cells were cultured for 24 h in RPMI 1640 medium supplemented with 10% FBS and 40 nM of Bortezomib (red coloring)(basal culture conditions for this cell line). The cells were then stained with the LIVE/DEAD kit and with Lysotracker RED. Equal numbers of events were acquired using a FACS Calibur flow cytometer by measuring the fluorescence of the LIVE/DEAD stain and Lysotracker RED, and the data were analyzed using the FlowJo software. The histograms display the median fluorescence intensity (MFI) of the Lysotracker RED channel. The figure presents a representative experiment. The same procedure was replicated three times.

To further study the phenomenon, western analyses of key autophagy biomarkers and regulators were performed, focusing on p62/SQSTM1, Atg5, Beclin-1 (Atg6), and LC3A/B (Atg8) (Figure 5a). p62/SQSTM1 is a cargo protein, transferring polyubiquitinated proteins for degradation through autophagy, thus linking the two degradational pathways (61). It interacts with LC3 II and the polyubiquitinated protein, leading to the protein’s degradation (62). During autophagy, p62 is degraded as well; therefore, the elevated p62 can indicate either autophagic flux suppression or increased autophagy, as it has been shown that p62 overexpression (as a result of NF-κB activation) serves the induction of autophagy (63). In the first case, other autophagic markers would be absent due to significant downregulation, while in the second case, increased p62 would be necessary to carry an increased load toward degradation. The role of p62 in Bortezomib resistance has only recently been uncovered, linking its expression with proteasome inhibitor-resistant cells (56). In our experiments, p62 was found to be significantly increased in the naïve cells after Bortezomib administration in a dose-dependent manner. In the resistant clone, regardless of the dose of Bortezomib applied, p62 remained relatively stable, with a baseline level three-fold greater than that of naïve cells (Figures 5a, 5g). This observation supports the exploitation of autophagy as an alternative proteostatic mechanism; however, other autophagic markers were assessed as well.

Atg5 was found elevated after incubation with Bortezomib in naïve cells and indicated a dose-dependent pattern of accumulation. The baseline accumulation of resistant cells is significantly greater than that of naïve cells. Incubation with 20 nM Bortezomib led to a slight downregulation in its accumulation, while the absence of Bortezomib or exceeding the baseline 40 nM dose led to its further accumulation (Figures 5a, 5h). Atg5 accumulation was considered an important marker indicating the upregulation of autophagy following both Bortezomib treatment and resistance emergence due to the cytoprotective nature of autophagy during stress conditions (64,65).

A protein of high significance in the study of autophagy is Beclin-1/Atg6, which regulates the autophagy-apoptosis axis through interaction with the Bcl-2 antiapoptotic protein family (66,67). Beclin-1 accumulation was observed to drop after treatment with a low dose of Bortezomib (20 nM; however, at the high, toxic dose of 40 nM, its accumulation increased, indicating the induction of autophagy due to severe UPS impairment. This was theorized to be the cell’s response to stress induced by Bortezomib, which activates autophagy to promote survival under these conditions. In the resistant cells, Beclin-1 maintains stable levels that are almost three-fold higher compared to the baseline expression in naïve cells. Sudden drug withdrawal or dose elevation seems to cause significant stress to the resistant cells as Beclin-1 levels rise even more (Figures 5a, 5i).

Finally, LC3A/B accumulation and conversion from LC3A/B I to LC3A/B II were assessed using western blots. Interpretation of LC3 accumulation comes with difficulties since its expression significantly increases during autophagy (LC3 I levels increase and, through conversion, LC3 II levels increase as well); however, the degradation of LC3 II inside the autophagosomes plays a significant role as well, thus decreasing detectable LC3 II. An accumulation of LC3 I can mean autophagic flux suppression, while an accumulation of LC3 II can be interpreted as autophagy induction (68).

Total LC3A/B was estimated as the sum of the two observed bands, one at ∼14 kDa (LC3 II) and one at ∼16 kDa (LC3 I). Following treatment with Bortezomib, the naïve cells exhibited a dose-dependent pattern of LC3 accumulation, because of the autophagic induction. LC3A/B increased more than two-fold following treatment with 40 nM of Bortezomib, while a clear upward trend was documented at the low dose of 20 nM (Figures 5a, 5j). The two forms of total LC3 were also analyzed separately (Figures 5k, 5l). In the naïve cells, LC3 I decreased by ∼40% following treatment with a low dose of Bortezomib (20 nM), while LC3 II increased its accumulation by ∼40%. At the high Bortezomib dose, both forms were found to accumulate more than two-fold compared to the untreated group. The ratio of LC3 conversion (LC3A/B II: LC3A/B I) which is commonly used to determine autophagy flux, exhibited a significant induction following the administration of 20 nM Bortezomib by almost 70% (indicating the UPS impairment), while the high dose returned the ratio to baseline levels (Table 2).

**Table 2.**
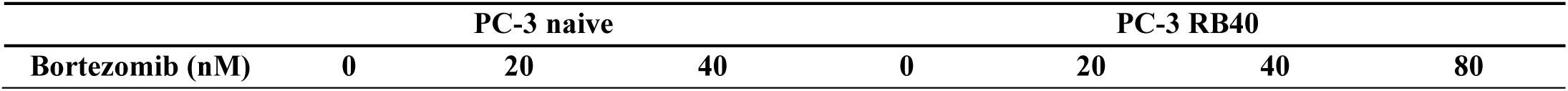
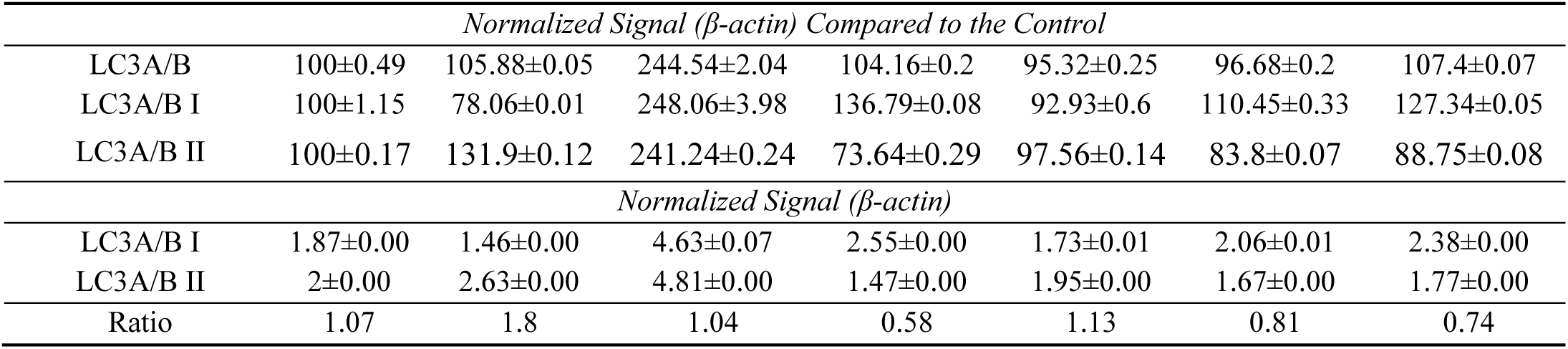
LC3A/B percentages. Tables should be placed in the main text near the first time they are cited.

Given the significant accumulation of both forms of LC3A/B, we did not theorize this restoration as an autophagy downregulation but rather an increase in degradation capacity that led to rapid LC3 degradation before its levels exceeded a particular detectable level. In the resistant cells, LC3 A/B fluctuated to slightly higher values compared to naïve cells. The baseline accumulation of total LC3A/B (40 nM of Bortezomib) was slightly lower than that documented in naïve cells. Deviations from this dose (total absence or higher drug concentrations) resulted in slight increases in LC3A/B accumulation, while the administration of a lower dose slightly decreased LC3A/B. A separate analysis of LCA/B I indicated that its presence followed the total protein accumulation pattern, while the results differed for LC3A/B II. LC3A/B II in general followed the same pattern as total LC3; however, at the 20 nM bortezomib dose, the resistant cells did not diminish their LC3 II levels but rather increased them. This could mean a rapid acceleration of autophagic flux; however, this would be controversial regarding the lesser interference imposed by a lower Bortezomib dose. Therefore, the elevation of LC3A/B II was interpreted using the accumulation of p62 as an index (since they are believed to directly interact (62)), leading us to the conclusion that when Bortezomib dose was diminished, the degradation capacity was tuned accordingly, by downregulating p62, and as a result, the accumulation of p62 increased. This interpretation also explained why the ratios calculated in resistant cells were lower compared to the control values, while all other autophagy indices assessed (Lysotracker, Atg5, Beclin-1, p62) supported the notion of upregulated autophagy. With autophagy being the sole targeted-degradation mechanism given that the UPS function was compromised, the autophagic degradation rate increased.

#### 2.2.3. Bortezomib-Resistant Cells Significantly Reduce their Intracellular Reactive Oxygen Species and Downregulate Stress Markers

The resistant cells’ stress levels were also assessed, focusing on the main stress markers and reactive oxygen species levels. The molecular chaperon Hsp70 is a protein that assists in UPS-mediated degradation and its expression in stress conditions. Bortezomib is a known Hsp family inducer, and this was documented in the naïve clone (27,69). Incubation with Bortezomib increased Hsp70 accumulation in a dose-dependent way (Figure 7a, 7b). This induction was not documented in the resistant clone. The PC-3 RB40 cells had baseline Hsp70 levels slightly greater than naïve PC-3 cells; however, its accumulation only showed signs of elevation after administration of 80 nM Bortezomib.

**Figure 7.**
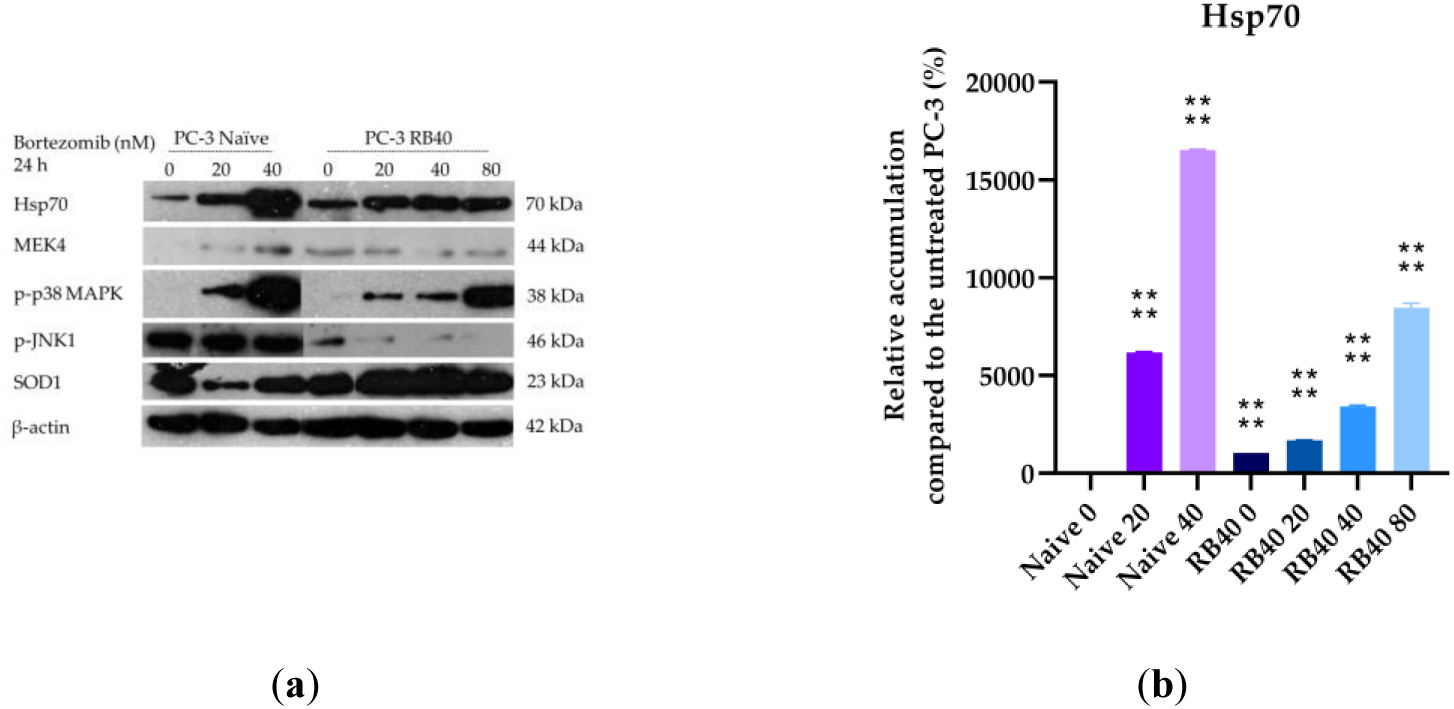
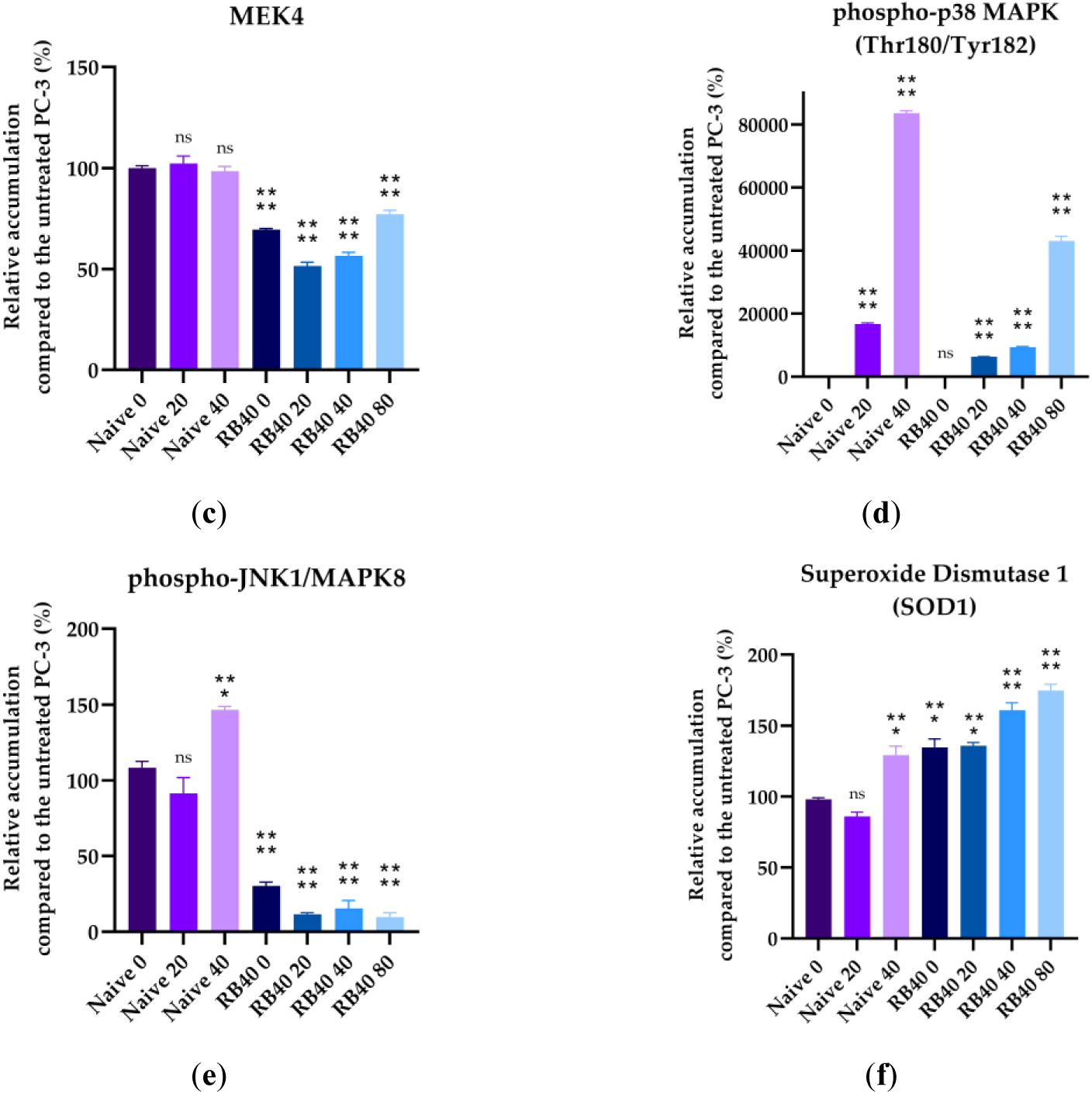
Western analysis of Hsp70, MEK4, p-p38 MAPK, p-JNK1, and SOD1. Cells (Naïve PC-3 and PC-3 RB40) were cultured inside 100 mm dishes, and 24 h before confluency, the media were changed, and fresh RPMI 1640 supplemented with 10% with or without the designated bortezomib doses (20, 40, 80 nM) was added. (**a**) The samples were prepared as previously noted, and by using specific polyclonal antibodies, the accumulation of Hsp70, MEK4, p-p38 MAPK, p-JNK1, and SOD1 was detected and developed on autoradiography films, which were subsequently scanned. Molecular weight markers were used during SDS-PAGE, and the approximate molecular weight of each detected polypeptide is annotated next to the band. (**b-f**) Quantification of the scanned blots was followed using the plug-in “Gel Blots” in ImageJ after conversion to grayscale images, and the band intensities and bit depth were calculated. The data were retrieved from triplicate experiments and after normalization using β-actin, bar charts were created. Each bar represents the average relative accumulation of the target protein compared to the untreated naïve PC-3 cells (first blot lane). The results were analyzed using multiple comparisons of one-way ANOVA in the Prism 8 software. (* corresponds to P =0,01; ** corresponds to P =0,001; *** corresponds to P =0,0001; and **** corresponds to P <0,0001). The error bars represent the standard error of the mean (SEM).

Other stress indicators that get activated after drug-induced stress are the MAPKs; p38 and JNK1 (70–74). Both MAPKs can be phosphorylated by MEK4, which was assessed as well to study the upstream regulation of the pathway. MEK4 was found to be significantly downregulated in the resistant clone, both during drug absence and in the presence of various doses. The highest Bortezomib concentration administered (80 nM) did not manage to raise MEK4 accumulation to naïve-like levels, while in the naïve cells, Bortezomib treatment resulted in MEK4 increasing (Figures 7a, 7c). p38 was assessed and found to be significantly phosphorylated after Bortezomib treatment in naïve cells following a dose-response manner, while the magnitude of the phenomenon was documented in the resistant cells and was many times smaller. In general, the resistant cells maintained low but detectable levels of p-p38 MAPK; however, the phosphorylation observed in the naïve clone was far greater even than those seen during the treatment of RB40 cells with 80 nM of Bortezomib (Figures 7a, 7d). The other major stress-related MAPK, JNK1, was assessed and found to follow a similar downregulation pattern in the resistant cells. In the naïve cells, upon treatment with 20 nM of Bortezomib, p-JNK1 initially decreased, while dose elevation to 40 nM led to a 50% increase compared to the untreated clone. The resistant cells were documented to have decreased JNK1 phosphorylation by 90%, a characteristic quite stable at different doses and drug absence (Figures 7a, 7e). These results indicated that Bortezomib, despite its action that renders PSMB5 non-functional upon binding, cannot successfully induce the signal transduction in stress-related pathways in the resistant cells, as they maintain low levels of the kinases (or the phosphorylated form of them) that can lead to apoptosis.

Given the importance of proteostasis in cell metabolism (through amino acid recycling, damaged protein degradation, and the role of ubiquitination as a way to regulate certain pathways), a direct link between proteasome function, autophagic flux, and intracellular oxidative stress was speculated. Therefore, the cells were assessed using H_2_DCFDA to monitor the generation of intracellular reactive oxygen species (Figure 8) (75,76).

**Figure 8.**
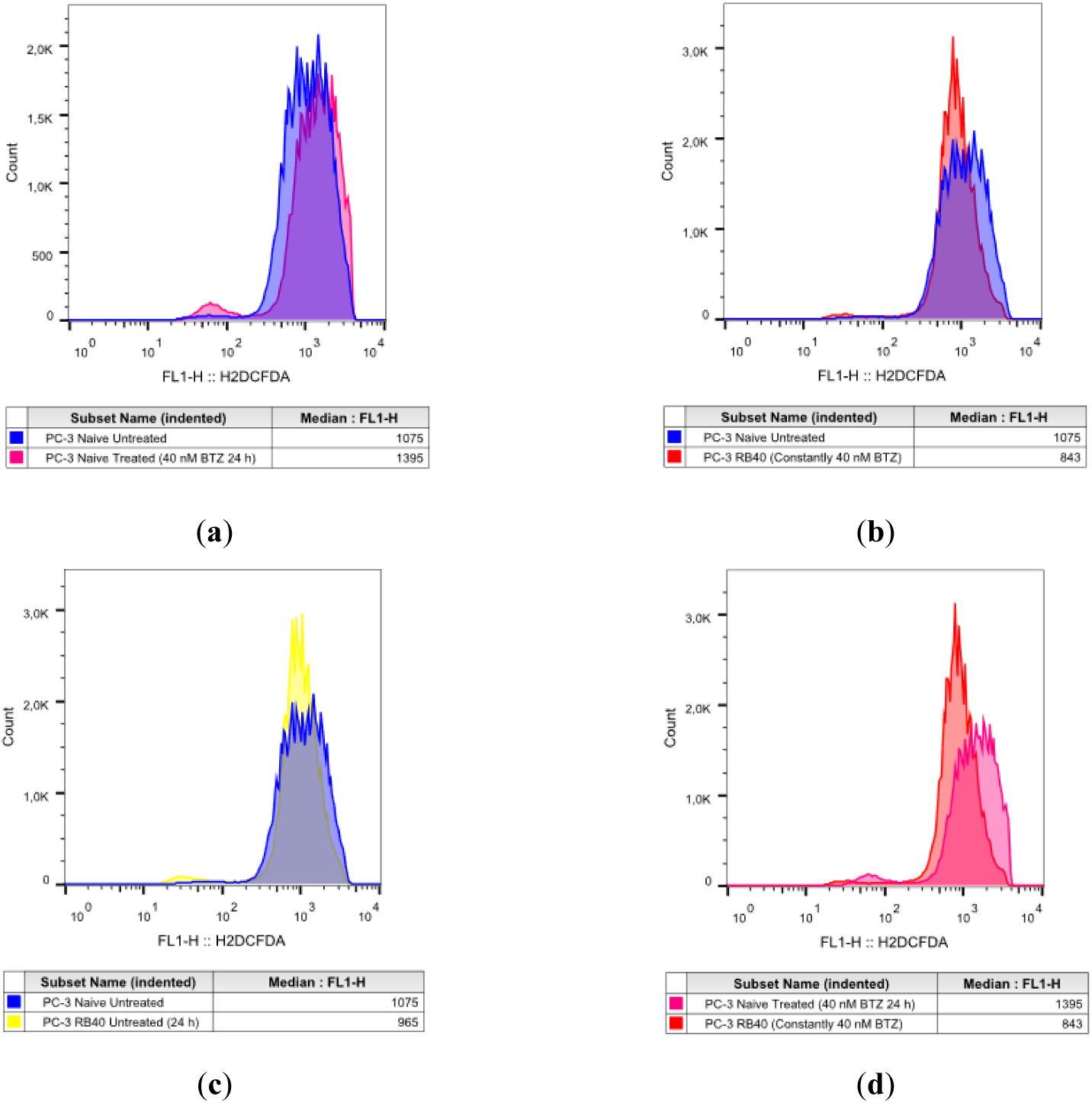
Intracellular Reactive Oxygen Species Assay of Naïve PC-3 and PC-3 RB40 Cells Following Treatment with Bortezomib. Equal numbers of cells were cultured inside 100-mm dishes, and 24 h before analysis, the media were replaced. (**a**) Naïve PC-3 cells were cultured for 24 h in RPMI 1640 medium supplemented with 10% FBS (blue coloring) (experimental control of baseline cell cycle progression); (**b**) Naïve PC-3 cells were cultured in medium containing 40 nM of Bortezomib for 24 h (magenta coloring); (**c**) PC-3 RB40 cells were cultured for 24 h in RPMI 1640 medium supplemented with 10% FBS (yellow coloring); and (**d**) PC-3 RB40 cells were cultured for 24 h in RPMI 1640 medium supplemented with 10% FBS and 40 nM of Bortezomib (red coloring) (basal culture conditions for this cell line). The cells were then stained with the LIVE/DEAD kit and with H2DCFDA. Equal numbers of events were acquired using a FACS Calibur flow cytometer by measuring the fluorescence of the LIVE/DEAD stain and H2DCFDA, and the data were analyzed using the FlowJo software. The histograms display the median fluorescence intensity (MFI) of the H2DCFDA channel. The figure presents a representative experiment. The same procedure was replicated three times.

Resistant cells were found to have lower oxidative stress levels compared to naïve cells, even after Bortezomib treatment (Figure 8b). This observation was made again by our research team in Zafeiropoulou et al. 2024 (27); however, it is validated here using another cell line and observing the same phenomenon. Time-course experiments showed that oxidative stress increases in both a dose-and time-dependent manner in naïve cells, while resistant cells always had significantly lower levels (Table 3).

**Table 3.**
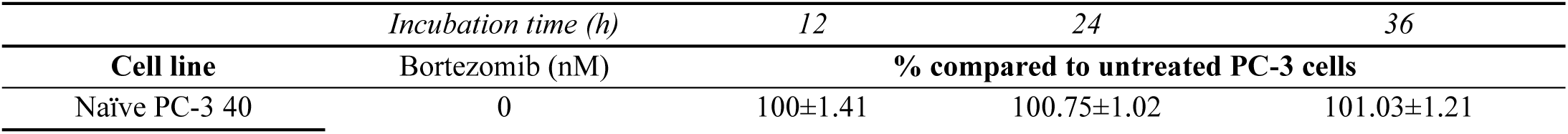
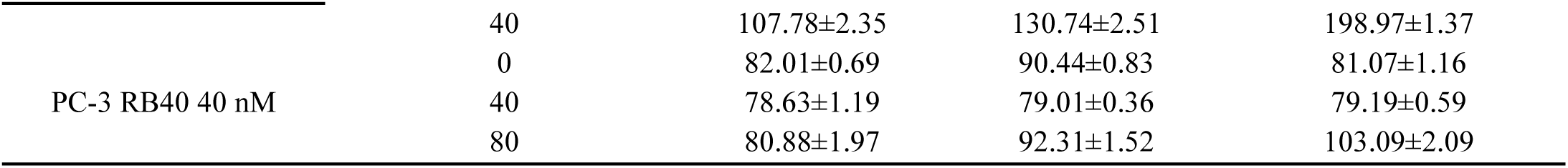
ROS levels percentages. Tables should be placed in the main text near the first time they are cited.

This was quite controversial given the impaired UPS and leads us to the conclusion that autophagy is responsible for recycling oxidatively damaged parts of the cells that normally would undergo K48 labeling and proteasomal degradation. The antioxidant enzyme superoxide dismutase 1 (SOD1) was also assessed due to its value as a marker (77). Increased SOD1 would indicate a constant need for superoxide radical degradation, as its synthesis is regulated by redox signaling, and would mean an increased antioxidant capacity of the cell. The RB40-resistant clone exhibited increased SOD1 levels, almost doubling its accumulation as shown in the blots, compared to the naïve clone, whose baseline SOD1 expression was lower (Figures 7a, 7f). Treatment with Bortezomib in both clones further increased the accumulation of SOD1, highlighting a correlation between redox signaling and Bortezomib that has not been fully uncovered.

### 2.3. Main Signal Transduction Kinases Found Upregulated and Cell Cycle Regulators Suppressed in Resistant Cells

#### 2.3.1. Bortezomib-Resistant cells Upregulate JAK/STAT, MAPKs, and PI3K/Akt Signaling Pathways to Evade Apoptosis

The main signaling pathways involved in Bortezomib resistance are the JAK-STAT, the Ras-Raf-MEK-MAPK, and the PI3K-Akt pathways (78–81). The main kinases were analyzed using Western analysis following 24 h of Bortezomib treatment with increasing doses. The cells were assessed at the 24-hour key point, as it was found to be the interval in which Bortezomib actions were most evident.

JAK is a kinase family that transmits signals from cytokines and growth factors to STATs. JAK1 was assessed and revealed to increase its accumulation at high Bortezomib doses both in naïve and resistant clones (Figures 9a, 9b). The lowest accumulation was documented both in naïve and resistant cells following treatment with 20 nM of Bortezomib. The absence of Bortezomib led to a double accumulation in the PC-3 RB40 cells, with almost identical accumulation levels documented at the 40 and 80 nM doses.

**Figure 9.**
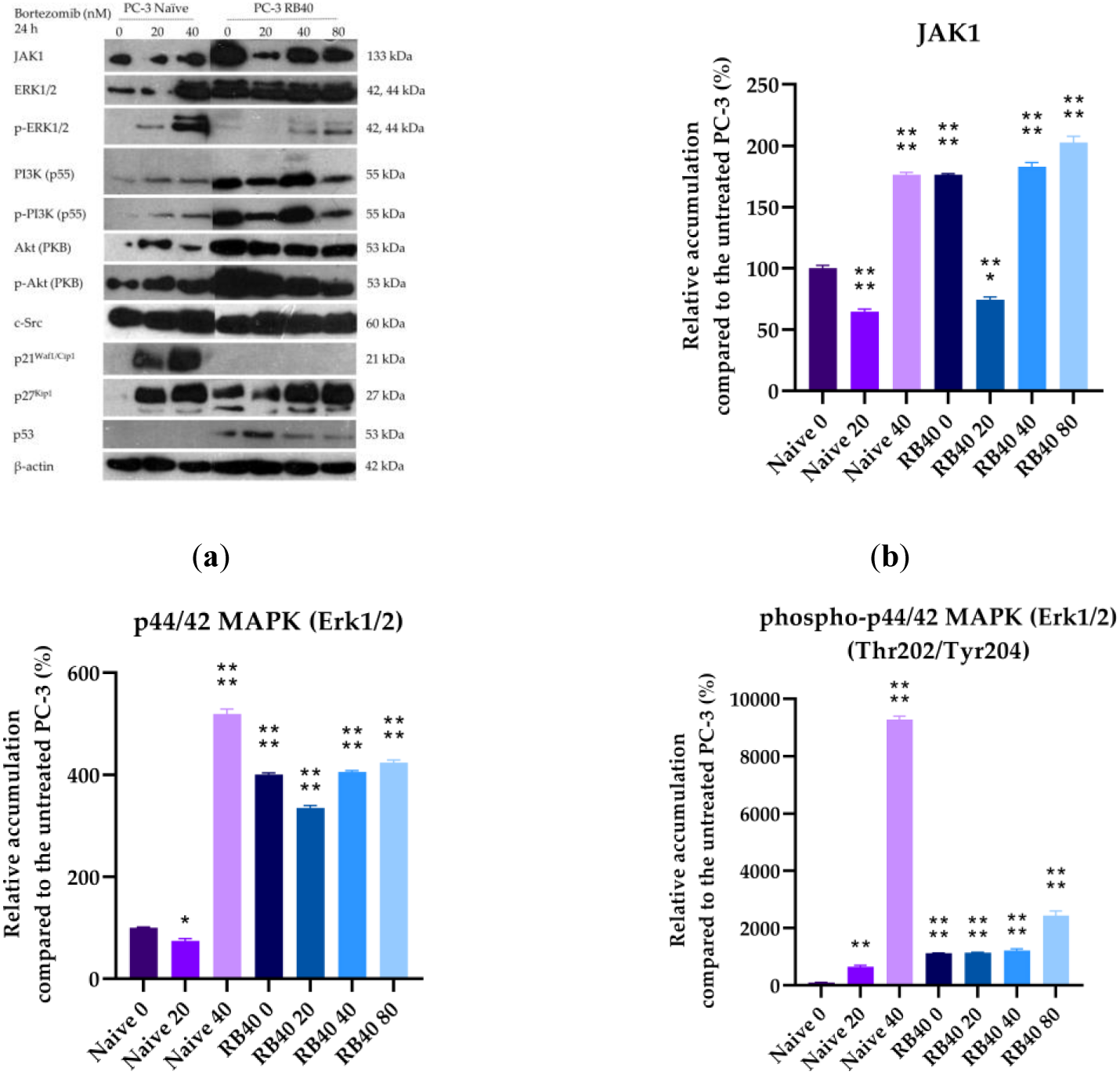
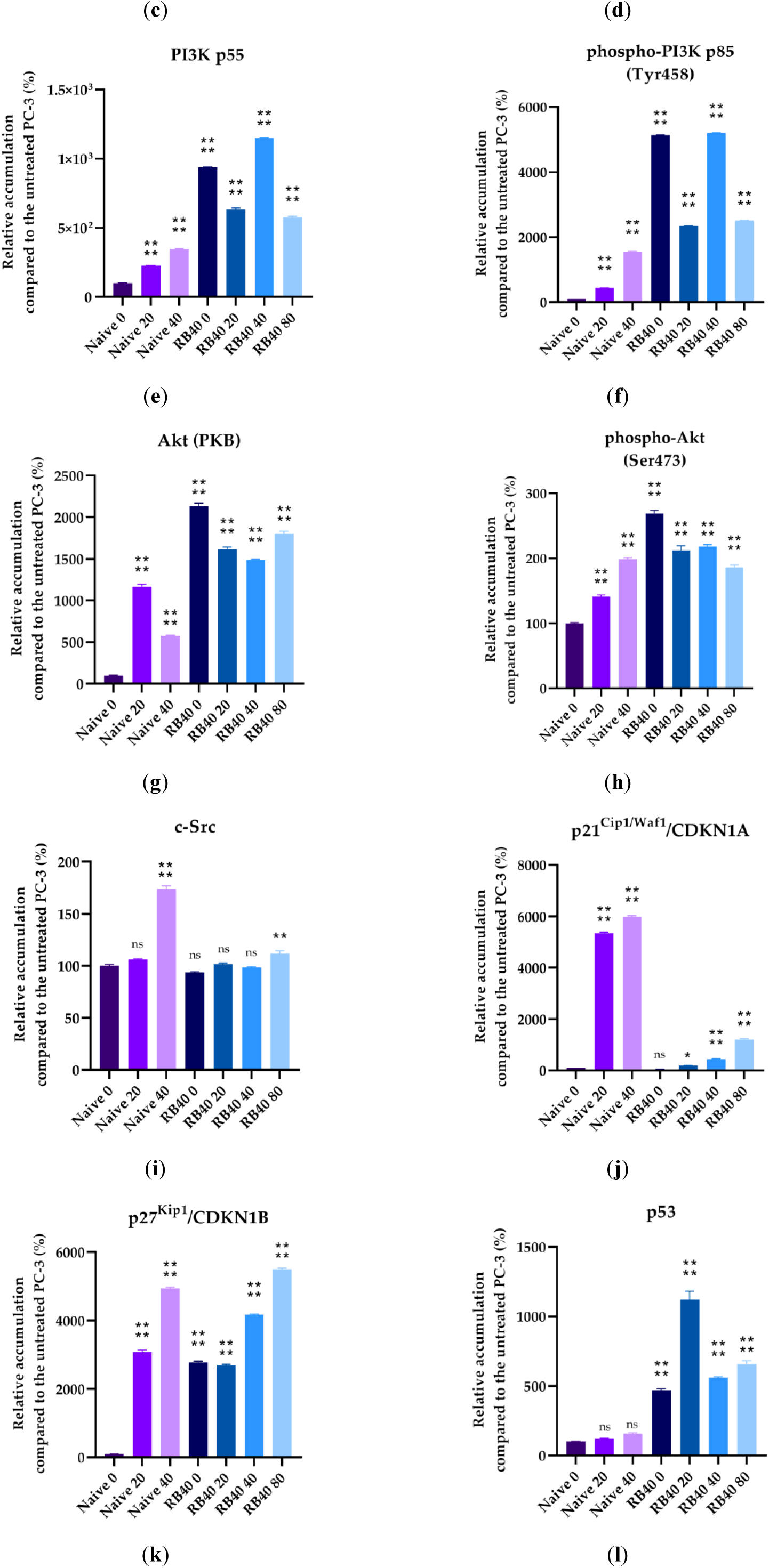
Western analysis of JAK1, ERK1/2, p-ERK1/2, PI3K, p-PI3K, Akt, p-Akt, c-Src, p21, p27, and p53. Cells (Naïve PC-3 and PC-3 RB40) were cultured inside 100 mm dishes, and 24 h before confluency, the media were changed and fresh RPMI 1640 supplemented with 10% with or without the designated bortezomib doses (20, 40, 80 nM) was added. (**a**) The samples were prepared as previously noted, and by using specific polyclonal antibodies the accumulation of JAK1, ERK1/2, p-ERK1/2, PI3K, p-PI3K, Akt, p-Akt, c-Src, p21, p27, and p53 was detected and developed on autoradiography films, which were subsequently scanned. Molecular weight markers were used during SDS-PAGE and the approximate molecular weight of each detected polypeptide is annotated next to the band. (**b-l**) Quantification of the scanned blots was followed using the plug-in “Gels” in ImageJ after conversion to grayscale images, and the band intensities and bit depth were calculated. The data were retrieved from triplicate experiments and after normalization using β-actin, bar charts were created. Each bar represents the average relative accumulation of the target protein compared to the untreated naïve PC-3 cells (first blot lane). The results were analyzed using multiple comparisons of one-way ANOVA in the Prism 8 software. (* corresponds to P =0,01; ** corresponds to P =0,001; *** corresponds to P =0,0001; and **** corresponds to P <0,0001). The error bars represent the standard error of the mean (SEM).

Regarding the Ras-Raf-MEK-MAPK pathway, we assessed the accumulation of ERK1/2, the foremost downstream kinases that can enter the nucleus and alter gene expression in favor of cell survival, proliferation, and regulation of apoptosis. The accumulation of ERK1/2 exhibited a five-fold increase in naïve cells after treatment with 40 nM Bortezomib, while lower doses (20 nM) did not alter ERK1/2 levels (Figures 9a, 9c). In the resistant cells, the baseline levels of ERK1/2 were four-fold higher compared to the control; only some fluctuation of minor significance was observed, with a decrease in ERK1/2 accumulation after administration of 20 nM Bortezomib and a slight elevation upon treatment with 80 nM of the drug. The phosphorylation patterns differed from those of the total proteins, as the resistant cells indicated a ten-fold change in ERK1/2 phosphorylation (Figures 9a, 9d). The naïve cells increased their p-ERK levels 90 times upon treatment with a high dose of Bortezomib, while a low dose increased the phosphorylation levels only six times. In the resistant cells, the 80 nM bortezomib dose increased the (already high) phosphorylation levels, doubling the amount observed in baseline conditions (40 nM).

Additionally, we assessed activity in the PI3K/Akt pathway by monitoring the two kinases’ accumulation and phosphorylation. This pathway regulates and promotes cell survival, induces autophagy as an anti-apoptotic mechanism, and transmits signals that come from upstream kinases, adhesion molecules, and other receptors (82). PI3K was found to accumulate in naïve cells, following a dose-dependent pattern of accumulation (Figures 9a, 9e). The resistant cells exhibited increased PI3K levels, at baseline Bortezomib levels (40 nM) which were maintained relatively stable. A reduction was observed in the low dose of 20 nM; however, during drug absence or high Bortezomib concentrations, the accumulation of PI3K was amplified. The phosphorylation patterns were analogous to total accumulation (Figures 9a, 9f). The resistant cells had activated PI3K, and its phosphorylation peaked at the 40 nM dose, indicating activation of the pathway to mediate the appropriate tuning of survival pathways. Downstream of PI3K, the protein kinase B (PKB), or Akt, was also assessed.

In prostate cancer, Akt activation is a marker of poor prognosis (83), and the PC-3 cell line was selected as a model of already defective Akt signaling since the phosphatase PTEN that suppresses Akt activation is absent in these cells, due to a double deletion (84). This lack of the tumor-suppressor gene PTEN makes PC-3 highly aggressive and more eager to develop drug resistance, as the presence of PTEN causes cell cycle arrest and subsequently apoptosis (85). Akt was found to be significantly upregulated in the resistant cells, fortifying the notion that the phosphorylation events already documented as well as the accumulation of other kinases may be transmitted through Akt (Figures 9a, 9g). Following treatment with Bortezomib, the naïve cells initially increase Akt accumulation (20 nM of Bortezomib), while greater doses reduce the initial augmentation. In the resistant cells, the baseline levels are higher than those observed in naïve cells, and deviation from the concentration that the cells are adapted to (40 nM), causes Akt levels to increase. This was interpreted as the cells’ response to the stress imposed by the new drug concentration that dysregulates the achieved equilibrium. To monitor the activation of Akt, we also assessed the accumulation of p-Akt which also validated the increased usage of the pathway (Figures 9a, 9h). Upon treatment with Bortezomib, the naïve cells increased the phosphorylation of Akt in a dose-response manner. The resistant cells exhibited a double accumulation of p-Akt compared to the naïve cells, and its accumulation slightly decreased as the drug dose increased. Since Akt is a downstream molecule and substrate of many intracellular kinases, it became evident that it acts as the other main signal transducer (with the second being the pair of ERK1/2) that transfers the appropriate messages inside the nucleus regarding the proteostatic capacity and regulates gene expression and cell cycle progressions as Akt is capable of overcoming the G_1_/S and G_2_/M arrest that can be caused by Bortezomib. The levels of the kinase c-Src were also assessed and found to be similar between naïve and resistant cells (Figures 9a, 9i). The sole exception was a peak in c-Src accumulation observed in naïve cells during treatment with 40 nM of Bortezomib. However, such an effect was not observed in the resistant cells.

#### 2.3.2. p21, p27 Found Suppressed in the Resistant Cells and p53 Fails to Cause Cell Cycle Arrest

One of the main effects of Bortezomib is the cell cycle arrest in the G_1_/S and G_2_/M phases, which is mediated through the accumulation of cell cycle regulators (86). Other studies have highlighted the accumulation of p21 and p27 after Bortezomib treatment (20–40 nM), which was evident in the naïve clone studied in this study. The resistant clone downregulated the accumulation of p21 to baseline levels and only started accumulating it again when the dose escalated to 80 nM of the drug (Figures 9a, 9j). Regarding the p27 accumulation, it was accumulated after Bortezomib treatment in naïve cells, and to a lesser point, it was also accumulated in the resistant clone (Figures 9a, 9k). An important cell cycle regulator is the p53 protein, which acts as a tumor suppressor gene. p53 normally inhibits cell cycle progression when it is overexpressed or accumulated; however, this was not observed during this study. The naïve cells accumulated p53 after Bortezomib administration, which was four-fold lower than the p53 observed in the RB40 cells (Figures 9a, 9l). The presence of p53 could promote DNA repair and protect the genome from the genotoxic effects of Bortezomib; however, its accumulation did not cause cell cycle arrest.

### 2.4. Nf-kB, STAT1, STAT3, Elk1, and cJun: Transcription Factors Regulating Resistance in the PC-3 RB40 Cell Line

Following the determination of the various kinases and intermediate molecules, our attention was directed toward the main transcription factors whose expression, accumulation and phosphorylation could have been altered. Nf-κB is the main transcription factor that has already been linked with Bortezomib resistance (87–89). In the naïve PC-3 cells, treatment with Bortezomib induced the accumulation of Nf-κB protein that was accumulated in a dose-dependent manner, and its phosphorylation increased accordingly (Figures 10a, 10b). In the PC-3 RB40 cells, the accumulation in the absence of Bortezomib was comparable to that of naïve cells (although slightly lower); however, in the presence of Bortezomib, its accumulation rose to high levels. Regarding phosphorylation, in the resistant cells, during incubation with 40 nM (the baseline for these cells) and 80 nM Bortezomib, Nf-κB exhibited the most activation, while withdrawal or reduction of the drug dose led to decreases (Figures 10a, 10c). In general, the activated fraction of Nf-κB in the resistant cells was significantly lower than that of naïve cells; however, the total Nf-κB levels were circa two times higher compared to the naïve clone.

**Figure 10.**
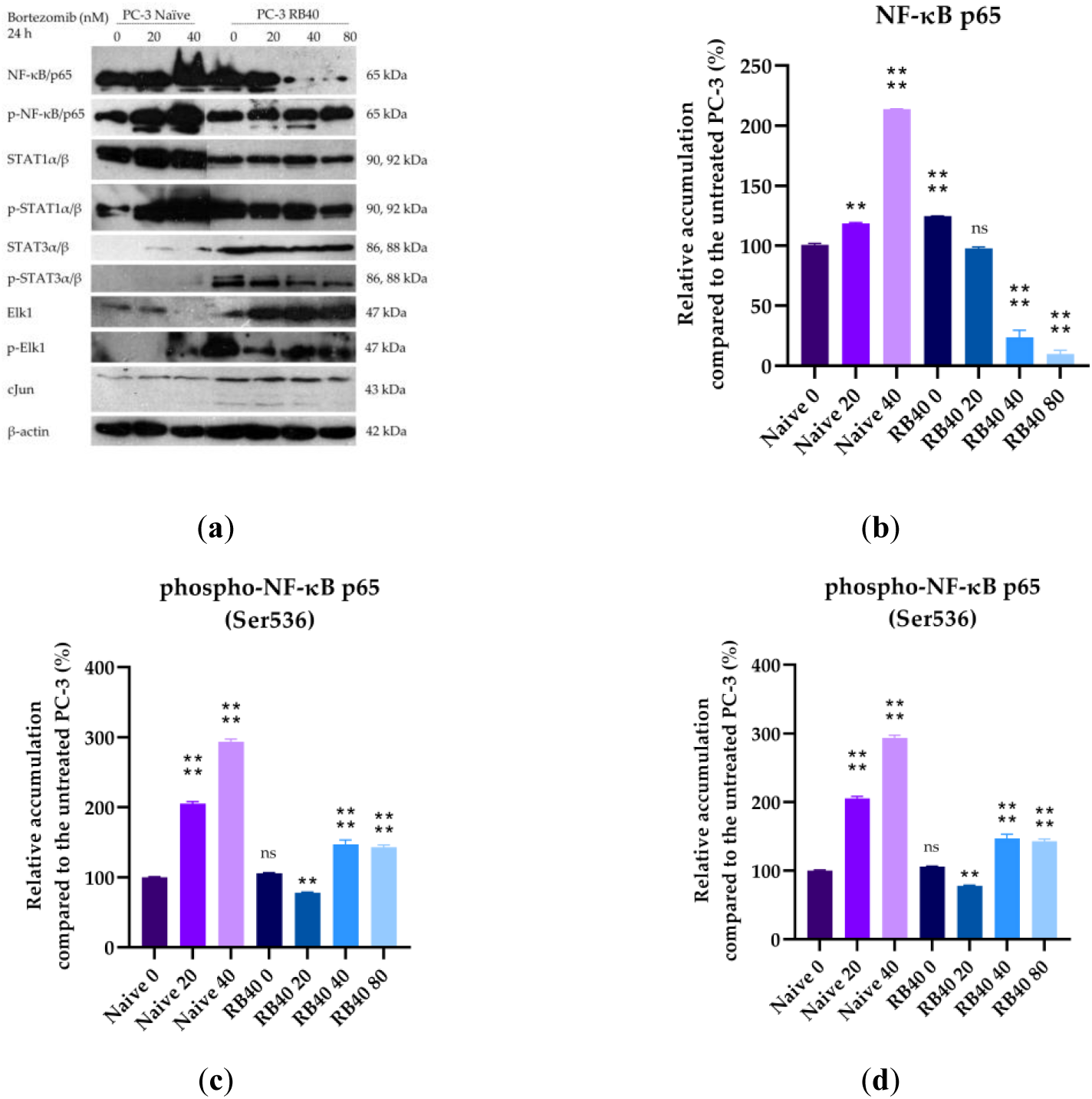
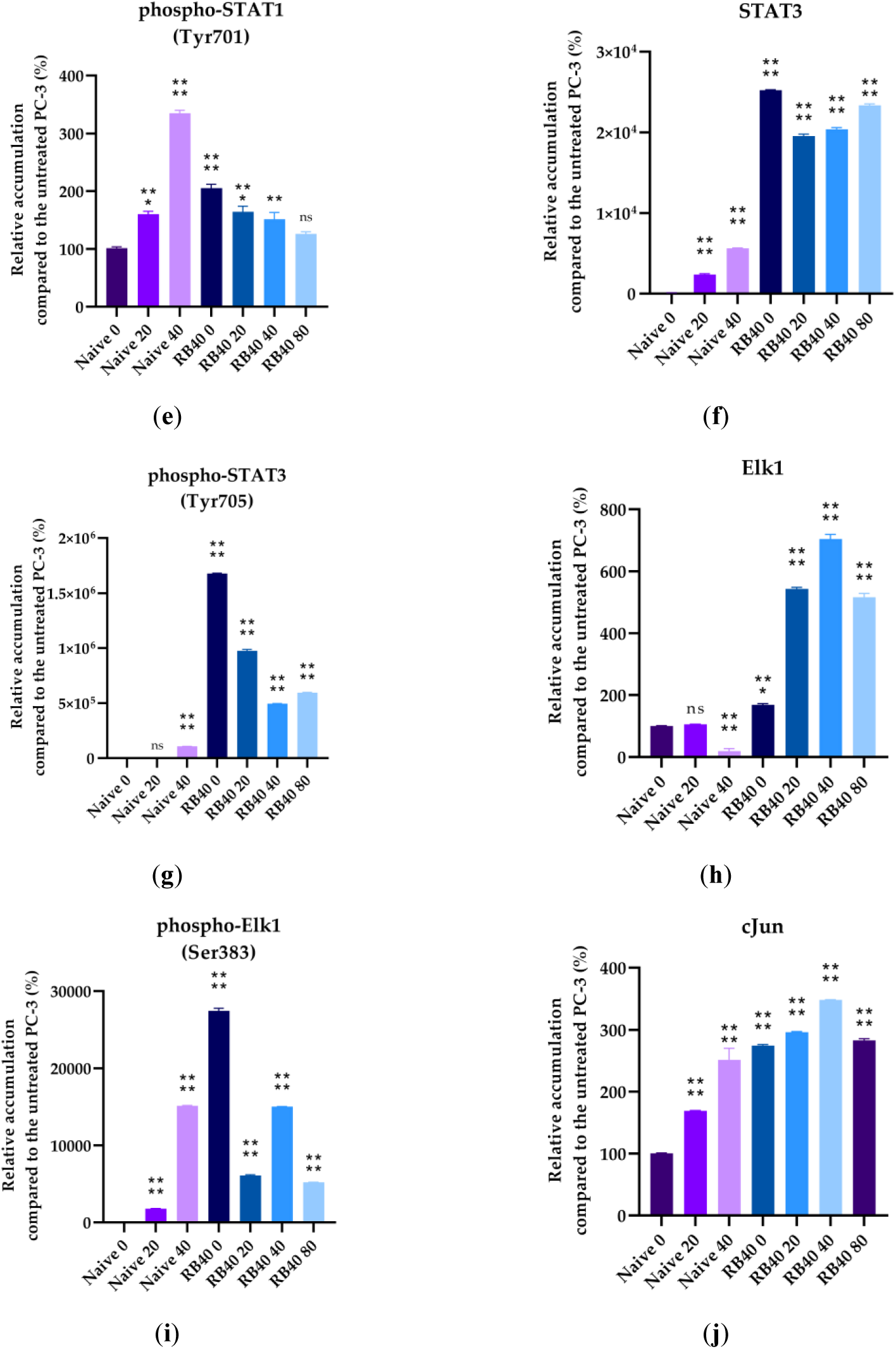
Western analysis of Nf-kB, p-Nf-κΒ, STAT1, p-STAT1, STAT3, p-STAT3, Elk1, p-Elk1, and cJun. Cells (Naïve PC-3 and PC-3 RB40) were cultured inside 100 mm dishes, and 24 h before confluency, the media were changed, and fresh RPMI 1640 supplemented with 10% with or without the designated bortezomib doses (20, 40, 80 nM) was added. (**a**) The samples were prepared as previously noted, and by using specific polyclonal antibodies the accumulation of Nf-kB, p-Nf-κΒ, STAT1, p-STAT1, STAT3, p-STAT3, Elk1, p-Elk1, and cJun was detected and developed on autoradiography films, which were subsequently scanned. Molecular weight markers were used during SDS-PAGE, and the approximate molecular weight of each detected polypeptide is annotated next to the band. (**b-j**) Quantification of the scanned blots was followed using the plug-in “Gel Blots” in ImageJ after conversion to grayscale images, and the band intensities and bit depth were calculated. The data were retrieved from triplicate experiments and after normalization using β-actin, bar charts were created. Each bar represents the average relative accumulation of the target protein compared to the untreated naïve PC-3 cells (first blot lane). The results were analyzed using multiple comparisons of one-way ANOVA in the Prism 8 software. (* corresponds to P =0,01; ** corresponds to P =0,001; *** corresponds to P =0,0001; and **** corresponds to P <0,0001). The error bars represent the standard error of the mean (SEM).

STATs are a group of transcription factors that are very important during cytokine signaling and are found to be activated in many cancer types. STAT1 and STAT3 are two members of the group with opposing biological actions, regarding the regulation of gene expression since STAT1 induces pro-apoptotic and anti-proliferative genes while STAT3 has pro-proliferative, anti-apoptotic, and pro-angiogenetic actions. Loss or downregulation of STAT1 is correlated with a poor prognosis (90), while induction of STAT3 activity also promotes the aggressiveness of the tumor (91). Regarding STAT1, incubation with Bortezomib led to a significant increase in its accumulation inside naïve cells that followed a dose-dependent pattern, acting as a precursor of apoptosis (Figures 10a, 10d). In the resistant cells, STAT1 was significantly downregulated (>50%). The phosphorylation levels of STAT1 rapidly arose in naïve cells following treatment with Bortezomib, in a dose-dependent manner, and reached a three-fold accumulation following treatment with 40 nM of Bortezomib (Figures 10a, 10e). In the resistant cells, the baseline levels of phosphorylated STAT1 were 50-80% upregulated compared to the naïve cells; however, the p-STAT1 levels never reached as high as those observed in treated naïve cells. STAT3 was found to be significantly elevated in the resistant clones compared to the naïve clone (which was not found at detectable levels) (Figures 10a, 10f). After treatment with Bortezomib, the naïve clone accumulated STAT3; however, this phenomenon was 1000-fold lower than that observed in resistant cells where the overexpression of STAT3 was evident and stable throughout different Bortezomib doses. Regarding STAT3 phosphorylation, the naïve cells mildly phosphorylated STAT3 after Bortezomib administration (compared to the untreated cells, where phosphorylation was undetectable), while the resistant cells constantly maintained fluctuating portions of phosphorylated STAT3 (Figures 10a, 10g). During drug absence (24 h), the resistant cells exhibited the highest levels of STAT3 phosphorylation, while the lowest values were observed at the basal conditions of 40 nM Bortezomib. Elevation of the dose increased the amount of phosphorylated STAT3, and so did the lower dose. Therefore, we concluded that the RB40 cells had adapted the 40 nM bortezomib presence, leading to a reduced need for STAT3 signaling compared to fluctuations in this dose which should be considered stress (both increases and decreases in the dose). In general, once compared to the naïve clone, the levels of phosphorylation in the PC-3 RB40 cells were two-to six-fold higher, highlighting the importance of STAT3 and its phosphorylation in achieving drug tolerance.

Elk-1 is a transcription factor whose role in Bortezomib resistance has not been thoroughly studied (92). Upon treatment with Bortezomib, the naïve PC-3 cells downregulated the accumulation of total Elk1 at the 40 nM dose, while Elk1 accumulation at the 20 nM dose remained relatively stable (Figures 10f, 10h). Regarding Elk1 phosphorylation, in naïve cells, the low dose of Bortezomib (20 nM) led to a 15-fold induction, while the high dose (40 nM) increased the abundance of the phosphorylated fraction even more (Figures 10a, 10i). In the resistant cells, Elk1 was observed in significantly greater quantities. In the absence of Bortezomib, the cells accumulated almost twice the amount of protein found in naïve cells, while incubation with Bortezomib significantly increased Elk1 presence. A peak in Elk1 accumulation was observed at the baseline dose of 40 nM, while fluctuations in this dose led to decreases in its abundance. The ratio of phosphorylated to total Elk1 remained stable during treatment with various Bortezomib doses, with peaks being observed in the absence of the drug and the usual dose of 40 nM.

cJun is a transcription factor activated in stress conditions, as a result of exposure to UV radiation, reactive oxygen species elevation, and drug-induced stress. Following Bortezomib incubation, cells have been shown to accumulate cJun which is believed to mediate apoptosis (74). Upon incubation with Bortezomib, the naïve cells accumulated cJun in a dose-dependent manner; however, the accumulation in the resistant clone was far greater. cJun was maintained at stable levels regardless of Bortezomib dose or even complete absence (Figures 10a, 10j).

## 3. Discussion

In this study, we created a Bortezomib-resistant prostate cancer cell line and proceeded to a broad-spectrum signaling assay by examining the main signal transduction pathways, transcription factors, and stress levels of the cells. Our model, the PC-3 cell line, has already been used as a Bortezomib-resistant cell line (93,94), as it can acquire resistance to proteasome inhibitors after prolonged exposure. Given the decreased effects Bortezomib has on many solid tumors, including prostate cancer, we tried to gather evidence on signaling molecules’ alterations as a way to improve our targeting against drug-resistant tumors and expand our biomarker repertoire. The resulting cell line was assessed for the restoration of its basic biological actions like cell viability, cell proliferation rate, the ability to surpass cell cycle arrest, and changes regarding the cell’s motility and metastatic potential. The resistant cells exhibited several EMT characteristics, with the most evident being cadherin switch and β-catenin upregulation. The resistant cells were able to downregulate the accumulation of p21, override the inhibitory actions of p27, and avoid the cell cycle arrest caused by p53. All three proteins are regulated through ubiquitination, and their turnover rate is heavily impaired in treated naïve cells, leading to significant cell cycle arrest in the G_1_ phase. However, the resistant cells were able to bypass those checkpoints and not exit the cell cycle, an effect that was considered the consequence of viability pathways’ activation. The PI3K-Akt pathway is the main survival pathway of the cells, and in the resistant clone, it was found to be significantly activated. The activation of Akt can be catalyzed by PI3K, which was found to be both overexpressed and activated in the resistant clone, as well as through crosstalk with other kinases. Akt is known to be able to override cell cycle arrest signals as well as enhance cytoprotective functions like autophagy (95–97). The same actions are believed to be mediated by the NF-κB pathways, which were also found activated in the resistant cells (98). NF-κB signaling is a known Bortezomib-resistance mechanism, initially observed in hematological malignancies where NF-κB is inherently active due to its participation in inflammatory reactions (28,87,89,99). In our case, the prostate cancer Bortezomib-resistant cell line PC-3 RB40 exhibited significant NF-κB activation and accumulation compared to the initial lower-level expression and activation of the pathway.

Our results indicated that the resistant clone created kept raising its tolerance against the drug the longer it was cultured in a Bortezomib-rich medium; however, no multidrug phenotype emerged. The mechanisms underlying Bortezomib resistance in the PC-3 RB40 cells were observed as the upregulation of PSMB5 synthesis, the utilization of autophagy as the main proteolytic pathway, and the suppression of oxidative stress levels to levels inferior to those observed in non-resistant cells. This observation was also made by our research grousing a Bortezomib-resistant DU-145 cell line (27). The resistant clone was able to downregulate the levels of poly-ubiquitinated proteins, an observation also made by using hepatocellular carcinoma and prostate cancer cell lines (93,100). In contrast to Yerlikaya and Okur (2019) where a Bortezomib-resistant cell line had also been created, we observed the mature PSMB5 overexpressed and no accumulation of the precursor form, indicting a different pathway of resistance (93). Recently, the role of p62 in proteasome inhibitor resistance has gained interest, given the protein’s role as a connection molecule between ubiquitin-proteasome degradation and ubiquitin-dependent microautophagy (56,57,61). The resistant cells were found to overexpress p62, confirming data from previous studies that all the autophagy markers assessed indicated a strong indication of the pathway during resistance (27,56,57,101). Autophagy modulation as an alternative proteolytic mechanism could explain the cells’ ability to control the turnover of various kinases and cell cycle inhibitors that otherwise accumulate and lead the cell to apoptotic death. An important role in this process was attributed to Beclin-1, which was also overexpressed in resistant cells. Aside from the regulation of autophagy, it has been found to directly interact with the Bcl-2 anti-apoptotic protein family, thus acting cytoprotectively (35,66,67). The modulation of signaling molecules with known cytoprotective roles was evident in the resistant clone, mainly focusing on Hsp70 and the stress-related kinases p38 and JNK1 (70,71,94,102). All three proteins are known to accumulate in stress conditions, an observation also evident in naïve cells following treatment with Bortezomib (13,74,103). The accumulation of MEK4 was also observed to be significantly downregulated in the resistant cells, and given the role of MEK4 in phosphorylating both MAPKs (p38 and JNK1 (104,105)), it could potentially act as a target for cell sensitization to Bortezomib. The downregulation of the stress markers directed us to oxidative stress levels, which were found to be suppressed. Oxidative stress is believed to rise inside a cell as a result of metabolism modulation, the accumulation of free radicals due to antioxidant defense failure, and specific chemotherapies that can dysregulate these cell functions. Bortezomib is a known oxidative stress-inducing molecule (9); however, this action was not observed in the resistant cells, indicating changes regarding metabolism-and antioxidant-related protein expression. The fortification of the resistant cells’ antioxidant defense systems was verified by assessing SOD1 levels, which were found to be overexpressed in the resistant clones. The reactive oxygen species inside the resistant cells, as measured using H_2_DCFDA, were lower even than those of untreated naïve cells, indicating the new redox equilibrium the resistant cells maintain independently of drug presence or absence. This had also been observed in the DU-145 cell line in a previous publication by our research team (27).

Given the importance of the signal transduction pathways in regulating these parameters, we focused on the main signaling pathways and examined the accumulation and phosphorylation of main kinases that participate in survival, proliferation, and anti-apoptotic gene expression, namely the JAK-STAT, Ras-Raf-MEK-ERK, and PI3K-Akt pathways. The JAK kinases, JAK1 and JAK2, have been found to participate in the pathogenesis of multiple myeloma, a malignancy susceptible to Bortezomib that can develop resistance as well (106). JAK silencing has been found to increase cell susceptibility to NK cell-induced cell death in multiple myeloma because both kinases transmit survival signals to the nucleus through the phosphorylation of STATs (107). Indeed, the resistant cells exhibited increased accumulation of JAK1, indicating a way to multiply pro-survival signals. The transcription factors STAT1 and STAT3, downstream of JAK1, were also assessed to monitor the signal transduction route. STAT1, with known pro-apoptotic roles (33,90), was found to be downregulated in the resistant clone both in terms of total STAT1 as well as phosphorylated STAT1. The opposite was observed regarding STAT3, which was significantly upregulated in resistant cells. Vangala et al. (2014) documented a correlation between STAT3 phosphorylation and PSMB5 protein levels, indicating STAT3 as a transcription factor that controls proteasome function (79). STAT3 inhibition downregulated the expression of proteasome subunits, thus increasing the pro-apoptotic effects of Bortezomib. Yuan et al. (2023) also proposed STAT3 inhibition as a way to overcome Bortezomib resistance (30). Both studies in multiple myeloma reported high STAT3 expression and phosphorylation, which was also observed in the resistant PC-3 RB40. This result also verified Zafeiropoulou et al. (2024) where the same phenomenon was documented in a Bortezomib-resistant DU-145 prostate cancer cell line (27).

Another major role in Bortezomib resistance is played by the ERK1/2 kinases. Being the foremost downstream molecules of the Ras-Raf-MEK-ERK pathway, they capture and transmit to the nucleus signals from several upstream kinases as well as from other pathways due to crosstalk between them (104). ERK1/2 phosphorylation has already been proposed as a molecular target in myelodysplastic syndromes, where the MEK inhibitors U0126 and PD98059 successfully re-sensitized Bortezomib-resistant cells to the drug (108). In our experiments, we did not inhibit the signal transduction; however, by assessing the pathway activity we observed that ERK1/2 was both upregulated and over-phosphorylated in the resistant clones. Survival mediated by ERK1/2 in Bortezomib resistance has also been shown in other studies, and prostate cancer is one of them (27,109). ERK1/2 are believed to regulate many cellular processes, from the transcription of survival-related genes to autophagy and even apοptosis. An important downstream molecule of the ERK1/2 kinases is the transcription factor Elk1, a molecule whose role in Bortezomib resistance has only been poorly studied so far.

Few pieces of evidence regarding Elk1 in proteasome inhibitor resistance have been published up to this day; however, the interaction between ERK1/2-Elk1 and the proteasome has already been proven (110). A study about Carfilzomib sensitivity in mantle cell lymphoma, a hematological malignancy for which proteasome inhibitors are an approved therapy, investigated the role of Elk1 in proteasome capacity and inhibitor resistance (92). Elk1 was found to control the proteasome assembling process, regulated by POMP (proteasome maturation protein) expression, which is a molecular chaperone regulating the biosynthesis of proteasome subunits. In that study, the activation of Elk1 through the c-Met/MAPKs, led to increased POMP transcription, which in turn increased the assembly of proteasome and increased both proteasomal activity as well as resistance against Carfilzomib (92). In our study, the MAPKs were also found activated and the most interesting fact is that Elk1 was found significantly overexpressed and phosphorylated in the PC-3 RB40 resistant cell line. Elk1 has recently been uncovered to play important roles regarding the progression of many cancer types including the aggressiveness of pancreatic, and prostate cancer (111–113). Modulation of Elk1 activity seems to have anti-metastatic effects as shown by Lai et al. (2023) where Asiatic acid was administrated in PC-3 cells and by interfering with the protein interaction between Elk1 and MZF-1, migration was negatively affected (114).

Another interesting finding regarding stress-response signaling pathways was the increased accumulation of cJun in the resistant clones. cJun is the last molecule in the JNK signaling cascade and is overexpressed in aggressive cancer types. cJun overexpression has also been linked to constitutively phosphorylated ERK1/2, a fact that was also observed in the PC-3 RB40 cell clone (115). Given the oncogenic functions of cJun, the suppression of cell cycle regulators like p21, and the proliferative and angiogenic effects of its action, it seems to be a key-molecule in resistant cell survival. Therefore, its role and expression in Bortezomib-resistant cell types should be expanded.

## 4. Materials and Methods

### 4.1 Cell Culture

The PC3 (ATCC, Manassas, VT, USA) cell line was used as a human prostate carcinoma cell model. The cells were grown in RPMI 1640 medium supplemented with 10% Fetal Bovine Serum (FBS), 100 units/mL penicillin, and 100 μg/mL streptomycin and maintained at 5% CO2 and 100% humidity at 37 °C. Cell culture media (RPMI 1640 with stable glutamine, pyruvate, and NaHCO3) and cell culture-related reagents (fetal bovine serum, 0.25% trypsin solution in PBS, and penicillin/streptomycin) were purchased from Biowest (Nuaille, France). Cell culture dishes, microplates, and Transwell chambers were from Greiner Bio-One (Kremsmünster, Austria). Flow cytometry expendables and reagents were from BD Biosciences (New Jersey, USA). The proteasome inhibitor Bortezomib was purchased from Janssen (Velcade^®^, Brussels, Belgium), Carfilzomib was from Amgen Inc. (Kyprolis^®^, Zurich, Switzerland), and Doxorubicin was purchased from Pfizer (Adriblastina^®^, Athens, Greece).

### 4.2. Proliferation Assays

Equal cell numbers were seeded inside 48-well culture plates and left to attach and grow for 24 hours. After this interval, the medium was aspirated, and fresh medium with increasing concentrations of Bortezomib (0, 5, 10, 20, 50, 100, 150, and 200 nM) was added, and the cells were incubated for 72 h with the drug. Each Bortezomib concentration was administered in triplicate wells. Subsequently, the media were aspirated, and the adherent cells (alive) were fixed with 4% v/v formaldehyde in PBS for 15 min and then stained with 0.5% crystal violet in 25% methanol for 20 min.

Following gentle rinses with water, the plates were left to air-dry, and the retained crystal violet was extracted using a 30% acetic acid aqueous solution. Afterward, the optical density at 595 nm was measured. The same procedure was followed to calculate the Carfilzomib IC_50_ (using the same concentration range as with Bortezomib) and the Doxorubicin IC_50_, incubating the cells with concentrations ranging from 10 to 1000 nM.

### 4.3. Creation of the Bortezomib-resistant cell line PC-3 RB40

To create a cell line resistant to the proteasome inhibitor Bortezomib, the procedure described by Zafeiropoulou et al. was followed with slight modifications (27). The IC_50_ of non-resistant cells was calculated (designated as naïve PC-3 cells) following 72 hours of Bortezomib incubation, and half of this concentration was added to cell culture dishes (with 75% confluency). The cells were left to grow under constant drug presence, and the medium was replaced every 72 hours, constantly maintaining the same Bortezomib concentration (5 nM) for three passages (∼14 days). After the cells adapted to this concentration, the drug dose was changed to 10 nM and kept for two passages (∼14 days). The same procedure was repeated for the 15, 20, 25, and 30 nM milestones. Raising the inhibitors’ dose was not well-tolerated by the cells, and many days were required for the cells to divide. The cells needed circa three months to reach the 30 nM milestone; they were maintained for four weeks at this concentration, and after this interval, they were supplemented with 35 nM of the drug for six weeks. Finally, the dose was stabilized at 40 nM of Bortezomib, while cell growth remained impaired for another two months. After about five months of ever-increasing Bortezomib doses (from 0 to 40 nM) and two months of stable 40 nM Bortezomib presence, the cells were adapted to the drug dosage, and assays concerning cell viability, migration, apoptosis, autophagy, intracellular signaling, and oxidative stress were performed. The resulting cell clone, resistant to the proteasome inhibitor, was named PC-3 RB40 (Resistant-Bortezomib 40 nM). As a control, naïve PC-3 cells of the same passage that had been stored in liquid hydrogen vapors were thawed (1-2 passages before the assays) and used to screen for any differences between resistant and non-resistant cells.

### 4.4. Scratch Assay

The scratch test/wound healing assay was used to assess the wound healing rate of both naïve and resistant cells. Cells were seeded inside 6-well cell culture plates and left to grow until the formation of a cell monolayer. Using a sterilized P200 pipette tip, cruciform scratches were done, and the culture medium was aspirated. The cells were rinsed gently with a warm PBS solution, and subsequently, a fresh medium containing increasing concentrations of Bortezomib (0, 20, 40, and 80 nM) was added. Each Bortezomib dose was monitored in triplicate. Photographs were taken using a microscope-mounted camera. The microplate was photographed at the key intervals of 0, 24, 48, and 72 h. The wound healing rate was calculated using the Wound Healing plugin for ImageJ (116).

### 4.5. Transwell/Boyden Chambers

Migration and chemotaxis were assessed in Transwell/Boyden chambers with 8 μm filter pores (117). A serum-containing medium was added to the lower compartment (with or without Bortezomib), and 2×10^4^ cells (suspended in serum-free RPMI 1640) were added to the insert. The cells were left to migrate for 24 h, and then the filters were fixed using a 4% v/v formaldehyde in PBS solution. Cells from the upper side of the filters were scraped, and the remaining (migrated) cells were stained using a 0.5% crystal violet solution. The filters were photographed under a microscope, and the total cells on the filters were counted using the Cell Counter built-in tool for ImageJ.

### 4.6. Flow cytometry

The FACS Calibur (BD Biosciences, Franklin Lakes, NJ, USA) was used to assess cell viability, apoptosis, lysosomal activity, and intracellular reactive oxygen species levels. The cells were incubated for specific time intervals in a Bortezomib-containing medium, and after that, they were collected by trypsinization and centrifugation. The cell number was estimated using a Neubauer hemocytometer, and equal numbers of cells were used for the analyses. To assess apoptosis, cells were incubated in Bortezomib-containing medium for 12, 24, and 48 h and then harvested as previously described. Afterward, they were stained with propidium iodide (PI) and Annexin V-FITC (#556420, BD Biosciences, Franklin Lakes, NJ, USA) for 15 min at room temperature in the dark (118). To measure the lysosomal activity, cells were stained with Lysotracker RED (Invitrogen™, Waltham, MA, USA) diluted in serum-free RPMI 1640 medium at 37 °C for 30 min in the dark. To measure ROS, cells were stained with H_2_DCFDA (Sigma-Aldrich, Darmstadt, Germany) at 37 °C for 30 min following an already described procedure (75,76). To analyze only the viable cells, the cell viability kit LIVE/DEAD (Invitrogen™, Waltham, MA, USA) was used in all stainings, and the cells were appropriately gated. Each flow cytometry experiment was conducted in triplicate. The subsequent analysis was performed with the FlowJo V10 software (BD Biosciences, Franklin Lakes, NJ, USA).

### 4.7. Western Blots

Cells were cultured without drugs/inhibitors for 24 h and then incubated with the designated concentrations of Bortezomib or Doxorubicin for varying times. Subsequently, they were washed twice with an ice-cold PBS solution and lyzed using RIPA buffer (Thermo Fisher Scientific, Waltham, MA, USA). The extracts were aliquoted and kept at −24°C until the analysis. Total proteins were determined using the Bradford assay. Equal amounts of total proteins were mixed with Laemmli’s Sample Buffer 2X solution containing 5% β-ME, and the samples were denaturated at 95°C for 10 min. Proteins were separated using 12% polyacrylamide gels and transferred to an Immobilon-P membrane (Millipore, Burlington, MA, USA) for 30 min using Towbin’s transfer buffer in a semi-dry transfer system as described in Zafeiropoulou et al (27). The membrane was blocked in TBS containing 5% skimmed milk and 0.1% Tween-20 for 1 h at 37°C. Membranes were then incubated with primary antibodies (Table 1) diluted in blocking solution overnight at 4°C, under continuous agitation. All antibodies were either from Cell Signaling Technology (Danvers, MA, USA) (annotated as ‘CST’ in Table 4), or Santa Cruz Biotechnology (Dallas, TX, USA) (annotated as ‘sc’ in Table 4).

**Table 4.**
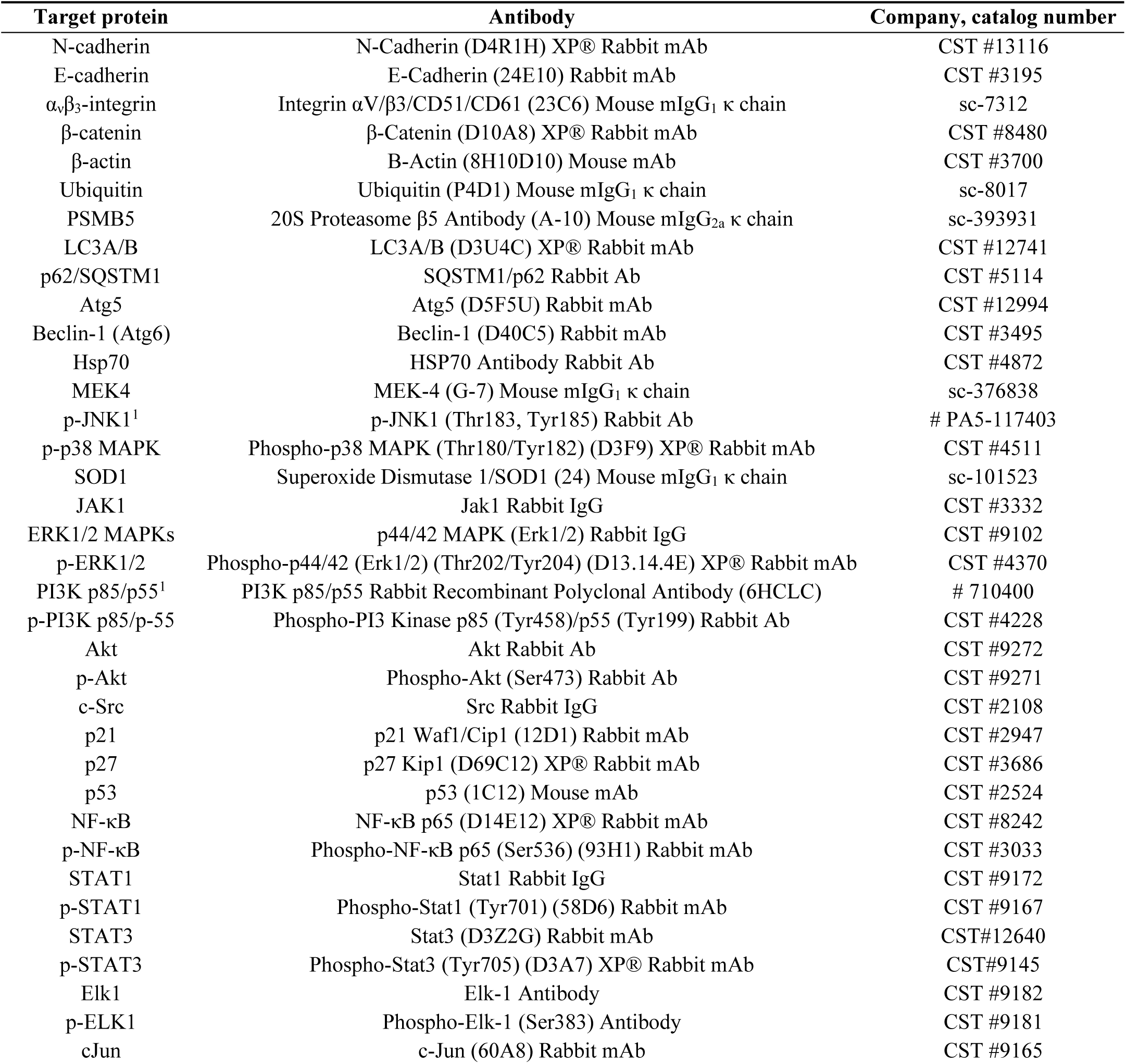
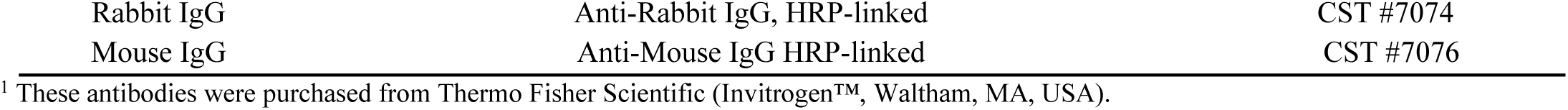
The antibodies used for western blot analysis.

The blot was then incubated with the appropriate secondary antibodies (Anti-rabbit IgG Antibody, CST#7074, or Anti-mouse IgG antibody, CST# 7076) (both diluted 1:2000) coupled to horseradish peroxidase, and bands were detected using the SuperSignal™ West Femto Maximum Sensitivity Substrate (Thermo Scientific™ #34096), according to the manufacturer’s instructions. Where indicated, blots were stripped in buffer containing 62.5 mM Tris HCl pH 6.8, 2% SDS, and 100 mM 2-mercaptoethanol for 30 min at 50°C and reprobed with primary antibodies. The blots were developed on autoradiography film (Fujifilm, Patras, Greece), and the films were scanned to analyze the band size and intensity. Quantitative estimation of the detected protein was performed through analysis of digital images using the ImageJ built-in ‘Gels’ tool. Each experiment was conducted three times and average intensities were extracted. Besides the Bradford assay which was used to calculate the loading volumes, the gels and membranes were stained with Coomassie Brilliant Blue after the analysis to validate the equal protein quantities. Actin was used as a reference protein since its accumulation did not exhibit fluctuation between naïve and resistant cells.

### 4.8. Statistical analysis

All statistical analyses were performed using Prism 8 (GraphPad, La Jolla, CF, USA) and Microsoft Office Excel. The proliferation assays were analyzed using Prism built-in non-linear regression equations to calculate the IC_50_ values. The wound healing rate was estimated using the average rate of three independent experiments as calculated using ImageJ. Flow cytometry data were analyzed in FlowJo and the differences between resistant and non-resistant cells were shown using t-tests in Prism 8. Western Blot data were collected using ImageJ and were subsequently analyzed using one-way ANOVA in Prism 8.

## 5. Conclusions

The model of our study, the PC-3 RB40 cell line, developed resistance to bortezomib, following exposure to gradually increasing doses of the drug for an extended time period. Our experiments demonstrated induction of autophagy, downregulation of oxidative stress levels, and increased signal transduction through the ERK1/2 and Akt and JAK kinases. Following investigation of various transcription factors, downstream of ERK1/2, Akt, we discovered that STAT1 was significantly downregulated in the resistant clone and STAT3 was both overexpressed and phosphorylated. Additionally, besides the activation of the Νf-κB pathway which is already extensively studied in cases of Bortezomib resistance, our study elucidated Elk1 and cJun as a potential pharmaceutical targets, given their elevated presence in the resistant clone. With the recent attention the MAPK-Elk1 axis has attracted and the evidence supporting STAT3 and cJun as potential therapeutic targets, we propose the three transcription factors as molecules of a significant role in the emergence of Bortezomib resistance, targeting of which could truly improve the management of prostate cancer cases, by expanding our drug repertoire and disease understanding.

## Author Contributions

Conceptualization, P.K. and G.K.; methodology, G.K., K.Z., T.A, S.A.; software, G.K., T.A., S.A.; validation, G.K., T.A..; formal analysis, G.K., T.A.; investigation, G.K., K.Z., T.A., S.A.; resources, A.S., P.K.; data curation, G.K. and T.A.; writing—original draft preparation, G.K.; writing—review and editing, A.S., P.K.; visualization, T.A.; supervision, P.K.; project administration, A.S., P.K.; funding acquisition, A.S. All authors have read and agreed to the published version of the manuscript.

## Data Availability Statement

The data used for this paper can be found within the paper and its supporting files.

## Acknowledgments

We kindly thank Ilias Kazanis (Department of Biology, University of Patras) for allowing us to operate the microscopy equipment of his laboratory, and Achilleas Theoharis (Department of Chemistry, University of Patras) for providing us with the western blot antibodies for STAT3 and phosphorylated STAT3.

## Appendices

## Appendix A

**Figure A1.**
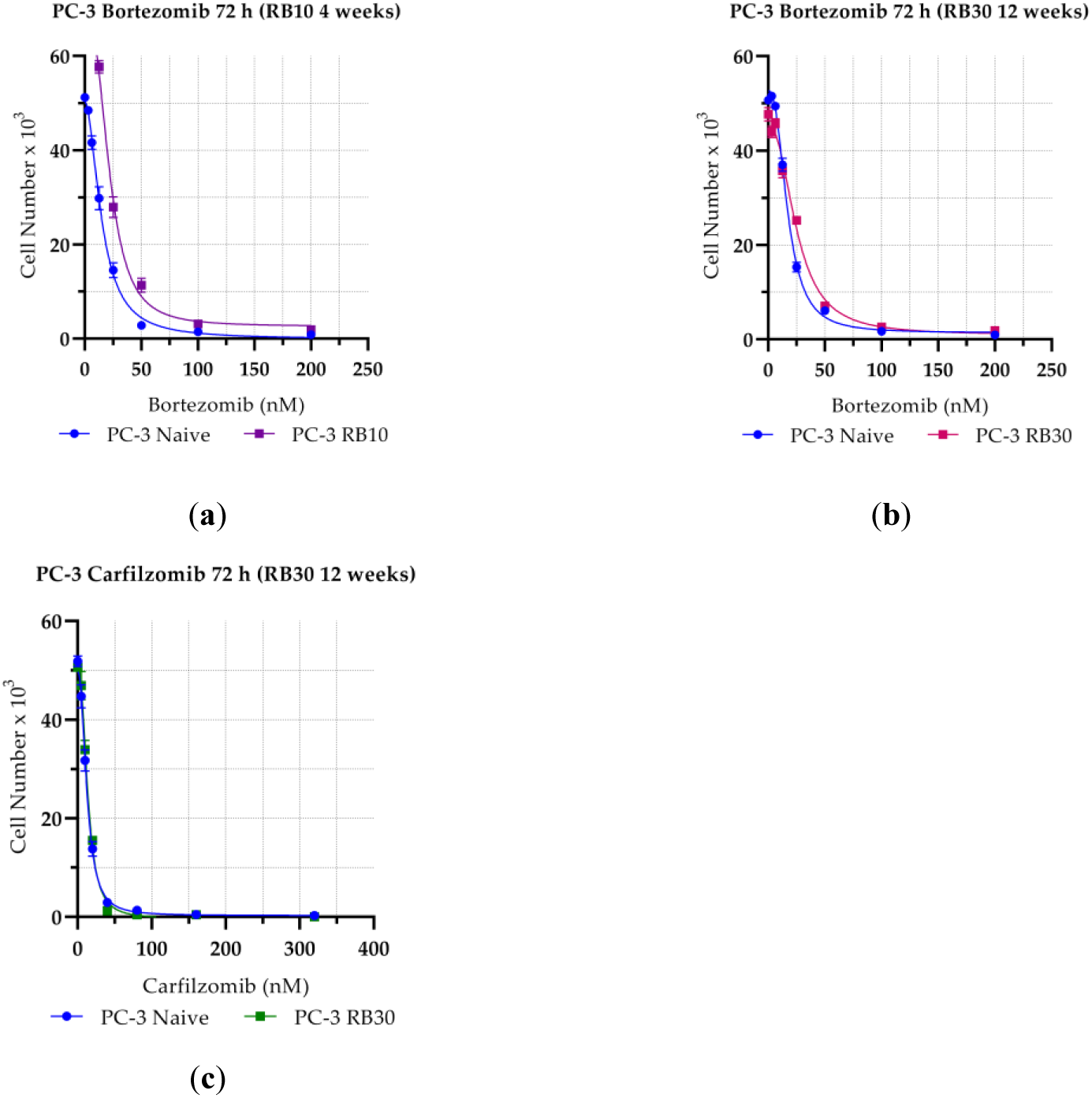
Assaying the Effects of Bortezomib and Carfilzomib on the Proliferation of Naïve PC-3 and PC-3 RB40 Cells During Dose Elevation. Equal numbers of cells were cultured for 72 h in 24-well microplates with various concentrations of (**a**) Bortezomib (0-240 nM) following 4 weeks of resistance acquirement; (**b**) Bortezomib (0-240 nM) following 12 weeks of resistance acquirement; and (**c**) Carfilzomib (0-325 nM) following 12 weeks of resistance acquirement; The live cells were measured using the Crystal Violet Assay. Each dot represents the average of three experimental values, and the error bars represent the standard error of the mean (SEM). The fitting line was graphed in Prism 8 using the built-in model for IC_50_ determination. Each plot represents one experiment, while the mean IC_50_ of three replicated experiments is presented in Table 1.

## Appendix B

**Figure A2.**
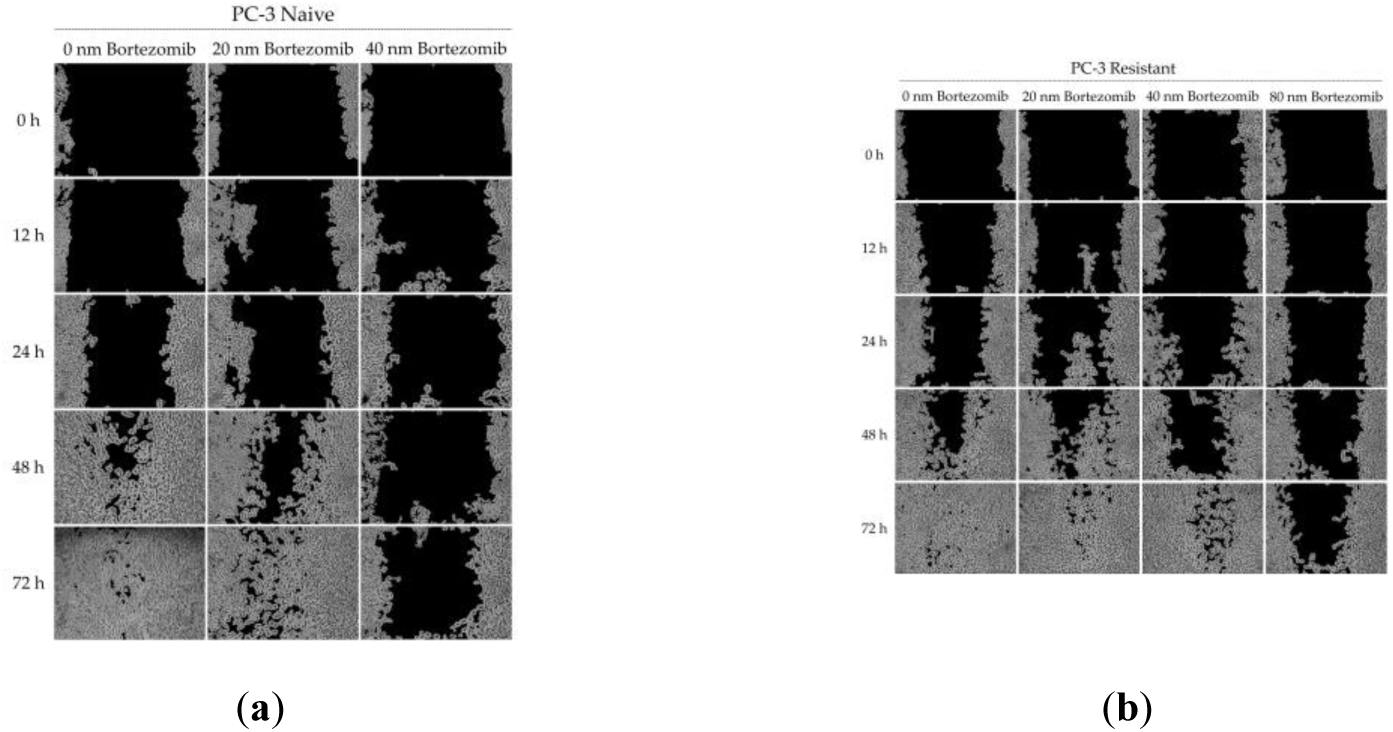
Scratch Test/Wound Healing Assay; Cells were cultured in 6-well plates until confluency, and then scratches were made. Various concentrations of Bortezomib (0, 20, 40, 80 nM) were added, and photographs were taken at the key time points of 0, 24, 48, and 72 h using a camera mounted on an inverted microscope at 100X magnification. The photographs showing wound closure were then analyzed using an ImageJ plug-in, and the healing rates were determined. The data were then plotted in Prism 8. (**a**) Naïve PC-3; and (**b**) PC-3 RB40.

**Figure A3.**
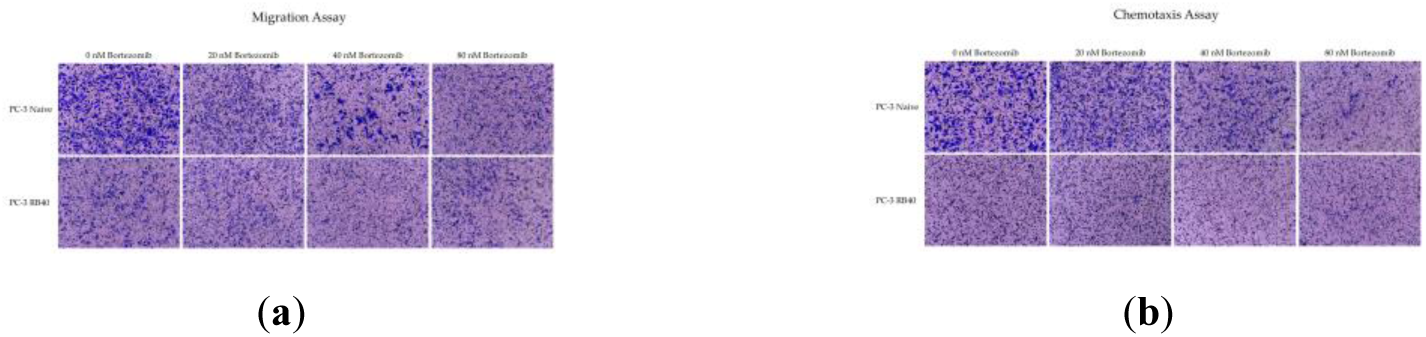
Boyden/Transwell Chambers. (**a**) Migration Assay; Equal numbers of cells were placed in Transwell/Boyden Chambers in serum-free RPMI 1640 medium containing increasing concentrations of Bortezomib (0, 20, 40, and 80 nM), and the inserts were placed in microwells containing FBS-supplemented medium. The cells were left for 24 h to migrate, and afterwards, the cells crossing the filters were fixed and stained with crystal violet. Photographs were taken using a 10x objective and the fixed cells were counted using the multipoint tool by ImageJ; (**b**) Chemotaxis Assay; Equal numbers of cells were placed in Transwell/Boyden Chambers in serum-free RPMI 1640 medium, and the inserts were placed in microwells with FBS-supplemented medium containing increasing concentrations of Bortezomib (0, 20, 40, and 80 nM). The cells were left for 24 h to migrate, and afterwards, the cells crossing the filters were fixed and stained with crystal violet. Photographs were taken using a 10x objective, and the fixed cells were counted using the multipoint tool by ImageJ.

